# Cell-type specific transcriptional adaptations of nucleus accumbens interneurons to amphetamine

**DOI:** 10.1101/2021.07.08.451674

**Authors:** David A. Gallegos, Melyssa Minto, Fang Liu, Mariah F. Hazlett, S. Aryana Yousefzadeh, Luke C. Bartelt, Anne E. West

## Abstract

Parvalbumin-expressing (PV+) interneurons of the nucleus accumbens (NAc) play an essential role in the addictive-like behaviors induced by psychostimulant exposure. To identify molecular mechanisms of PV+ neuron plasticity, we isolated interneuron nuclei from the NAc of male and female mice following acute or repeated exposure to amphetamine (AMPH) and sequenced for cell type-specific RNA expression and chromatin accessibility. AMPH regulated the transcription of hundreds of genes in PV+ interneurons, and this program was largely distinct from that regulated in other NAc GABAergic neurons. Chromatin accessibility at enhancers predicted cell-type specific gene regulation, identifying transcriptional mechanisms of differential AMPH responses. Finally, we observed dysregulation of multiple PV-specific, AMPH-regulated genes in an *Mecp2* mutant mouse strain that shows heightened behavioral sensitivity to psychostimulants, suggesting the functional importance of this transcriptional program. Together these data provide novel insight into the cell-type specific programs of transcriptional plasticity in NAc neurons that underlie addictive-like behaviors.

## Introduction

Drugs of abuse, including the psychostimulants amphetamine (AMPH) and cocaine, lead to addiction by driving progressive and lasting adaptations in the function of neurons within the mesolimbic dopamine reward circuit^1^. Psychostimulant-induced changes in gene transcription play an essential role in this process by persistently altering the functional connectivity of neurons in reward circuits^2^. These transcriptional responses can be accompanied by regulation of the epigenome, including dynamic modifications of histone proteins and direct modifications to genomic DNA^3, 4^. These biochemical marks may indicate fundamental changes to chromatin architecture in psychostimulant-activated neurons, or they could reflect the differential activation of pre-existing chromatin states^5^. Importantly, because chromatin architecture is highly cell-type specific, elucidating the relationship between chromatin regulation and gene transcription requires isolation and differential analysis of specific cell types from heterogeneous brain regions.

The Nucleus Accumbens (NAc) is a key region mediating the development and expression of addictive-like behaviors, and it is a major locus of psychostimulant-induced transcriptional and synaptic changes. The cellular and molecular consequences of psychostimulant-exposure have been well-documented in Spiny Projection Neurons (SPNs), which are the most numerous NAc neurons and provide the main output from this brain region. However, a growing number of studies suggest functions of local circuit interneurons of the NAc in the regulation of addictive-like behaviors. Despite comprising only a few percent of all NAc neurons, interneurons can exert dominant roles over SPN output^6^. For example, cholinergic interneurons are activated by cocaine and optogenetic suppression of this activity impairs cocaine-induced conditioned place preference^7^. By contrast, repeated cocaine exposure reduces the excitability of Somatostatin-expressing (SST+) GABAergic interneurons of the NAc, yet optogenetic activation and suppression of these neurons respectively enhanced and impaired cocaine-induced place preference^8^. Diverse consequences of altered interneuron activity likely arise from the ways these cells modulate SPN activity. However, the circuit-level mechanisms of NAc interneuron function are only beginning to be elucidated.

PV+ GABAergic interneurons are especially potent regulators of feed-forward inhibition in local striatal circuits, and experimental manipulations have implicated these neurons in long-lasting NAc circuit adaptations that promote addictive behaviors^9, 10^. PV+ interneurons robustly fire in response to psychostimulant exposure *in vivo*, and withdrawal after repeated cocaine exposure further increases their excitability^11, 12^. Excitatory inputs from basolateral amygdala to PV+ interneurons in the NAc shell are enhanced following cocaine self-administration, resulting in increased feedforward inhibition of NAc SPNs and more efficient encoding or training of the operant behavior^13^. Blocking neurotransmitter release from NAc PV+ interneurons prevents the expression of locomotor sensitization and conditioned place preference induced by repeated AMPH^14^. PV+ interneuron silencing leads to a global disinhibition of both D1 and D2 dopamine receptor expressing SPNs in the NAc, suggesting that blocking the inhibitory function of these interneurons may impair the expression of addictive-like behaviors by disrupting the SPN ensembles that encode the reward-related behavior^14, 15^.

Given the challenge of isolating interneuron populations from the brain for biochemical assays, little is known about psychostimulant-induced gene transcription and chromatin plasticity in interneurons. However, in our prior studies of the methyl-DNA binding protein MeCP2 we made the incidental discovery that psychostimulant drugs of abuse selectively induce MeCP2 phosphorylation at Ser421 in PV+ GABAergic interneurons of the NAc^16^. Furthermore, we found that transgenic mice bearing a phosphorylation site mutation (Ser421Ala) knocked into *Mecp2* both rendered them behaviorally supersensitive to psychostimulants and caused dysregulation of AMPH-dependent Fos expression in NAc PV interneurons^17, 18^. Here, to study both chromatin and gene expression regulation in NAc interneurons following AMPH exposure, we used the Isolation of Nuclei Tagged in Specific Cell Types (INTACT) transgenic mouse model^19^ to purify nuclei from PV+ or SST+ interneurons of the NAc. We identified hundreds of AMPH-regulated genes in both populations of interneurons and used differential patterns of chromatin accessibility to discover their mechanisms of cell-type specific regulation. Taken together these studies significantly expand our understanding of the transcriptional plasticity mechanisms that underlie the function of NAc interneurons in the neural response to psychostimulant drugs of abuse.

## Results

### Isolation of NAc interneuron nuclei enables cell type-specific sequencing

To isolate NAc interneurons for gene expression and chromatin accessibility measures, we genetically tagged the nuclei of specific neuronal cell-types using INTACT transgenic mice^19^ (**Fig. 1A-C**). When were crossed with *Pvalb*-IRES-Cre or *Sst*-IRES-Cre mouse lines, the nuclear Sun1-GFP transgene colocalized with PV protein (**Fig. 1D**) or *Sst* RNA (**Fig. 1E**).

**Figure 1:**
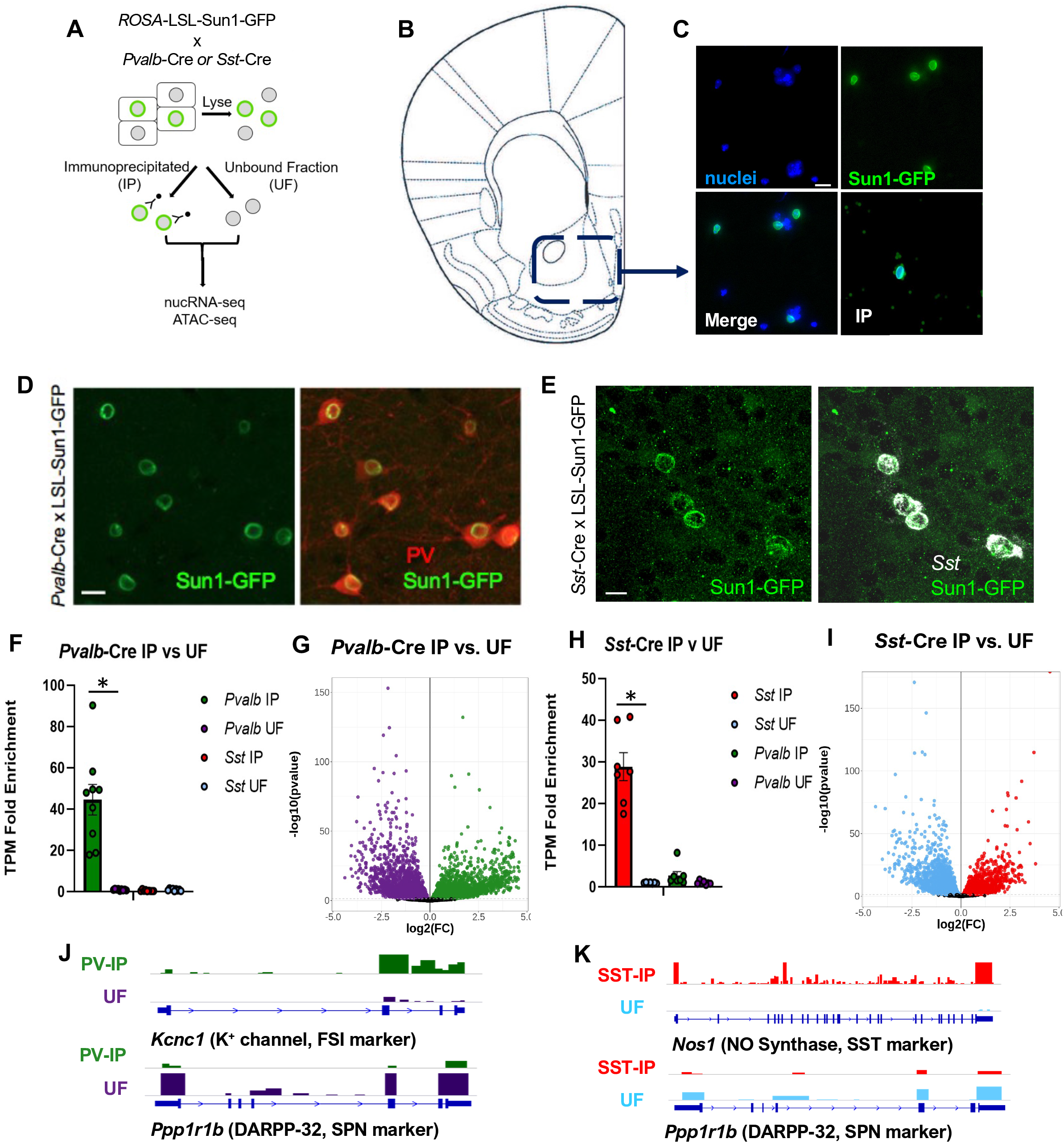
INTACT-mediated isolation of PV+ and SST+ GABAergic interneuron nuclei from mouse NAc. **A)** Schematic of the INTACT Cre-inducible Sun1-GFP transgene system. For each strain, the protocol yields two fractions: Sun1-GFP+ nuclei (green) immunoprecipitated from the specific cells expressing the Cre transgene (IP) and an unbound fraction (UF) that contains a mixture of nuclei from all other cell types present in the homogenate. RNA and chromatin from each fraction was used for nuclear RNA sequencing (nucRNA-seq) and the detection of Tn5-transposase accessible regions (ATAC-seq), respectively. **B)** Diagram of the NAc region bilaterally dissected from individual animals for nuclear isolation. **C)** Representative images of DAPI-stained nuclei and Sun1GFP-fluorescence in the NAc homogenate (merge), and on beads after immunoprecipitation (IP). **D-E)** Immunohistochemical overlap of Sun-GFP with Parvalbumin (PV) immunostaining (D), or *Sst* FISH signal (E), in coronal brain sections through the NAc of the indicated INTACT transgenic mice. Scale bars, 10µm. **F-I)** NucRNA-seq RNA expression data from IP and UF fractions of the indicated dual transgenic mice. *Pvalb*-Cre IP, green; *Pvalb-*Cre UF, purple; *Sst-*Cre IP, red; *Sst-*Cre UF blue; **F, H)** Validation in nucRNA-seq data of enrichment for cell-type marker genes *Pvalb* (F) and *Sst* (H) in the IP fraction of the indicated mice shown via TPM, Transcripts per Kilobase Million (TPM), Error bars indicate SEM. **G, I)** Volcano plots of cell-type enriched genes in *Pvalb*-Cre (**G**) and *Sst-*Cre (**I**) IP each compared to UF of the same strain in timepoint-combined control-treated mice. Black dots, not significant; colored dots, *FDR<0.05. *Pvalb*-Cre n=9, *Sst-*Cre n=7 individual animals. **J, K)** Representative nucRNA-seq tracks for cell-type marker genes. Y-axis is constant between matched samples for each gene. Arrows on gene indicate transcript directionality.

Immunoisolated GFP+ NAc nuclei from these mice (IP) showed enrichment for known cell-type markers compared to the unbound fraction (UF) by qPCR (**Fig S1A,B**) showing that this method is able to specifically isolate interneuron subtypes in NAc.

Most neurons in the NAc are GABAergic, with Spiny Projection Neurons (SPNs) representing the predominant cell type^20^. By contrast, each class of GABAergic interneuron comprises only a few percent of the total neurons^6^. To identify genes that are enriched in PV+ and SST+ interneurons, we performed RNA-seq on INTACT purified PV+ or SST+ nuclei from single mice (n=9 *Pvalb*-Cre, n=7 *Sst-*Cre) and identified genes that were differentially expressed relative to the UF from each respective pulldown. Although nuclei contain only a subset of total cellular RNA, prior studies have shown that nuclear RNA-seq (nucRNA-seq) gives a quantitatively accurate assessment of cellular gene expression that is robustly preserved upon dissociation of adult brain tissue^21^.

We identified 3145 genes including *Pvalb* (**Fig. 1F**) that were enriched in NAc PV+ interneurons nuclei relative to the UF of *Pvalb-*IRES-Cre mice, and 3108 genes that were de-enriched in the PV+ IP (**Fig. 1G; Table S1**). We identified 2273 genes including *Sst* (**Fig. 1H)** that were enriched in the SST+ IP nuclei relative to the *Sst-*IRES-Cre UF, and 2522 genes that were de-enriched in the SST+ IP (**Fig. 1I; Table S1)**. Among the IP-enriched genes, we found known markers of interneuron function, such as the voltage-gated potassium channel *Kcnc1* in PV+ neurons^22^ (**Fig. 1J**), the enzyme *Nos1* in SST+ neurons^23^ (**Fig. 1K)**. For both strains, the SPN marker *Ppp1r1b*, which encodes the signaling protein DARPP-32, was significantly lower in the IP nuclei relative to the fraction in the UF, consistent with the expectation that SPNs comprise the major fraction of cells found in the UF (**Fig. 1J, K).**

### AMPH induces a rapid program of transcription that overlaps between NAc neuron types

Neuronal activation induces multiple waves of stimulus-regulated gene transcription that can be separated by their timing and underlying mechanisms, including both a rapid and a delayed program of primary response genes (PRGs) driven by the post-translational modification of constitutively expressed transcription factors, as well as a delayed program of secondary response genes (SRGs) mediated by transcription factors (TFs) synthesized in the primary wave^24^. To identify rapid PRGs induced by AMPH exposure in NAc GABAergic interneurons, we purified PV+ or SST+ interneurons from the NAc of mice 35min following an injection of either saline as control or 3mg/kg AMPH in an open field chamber (**Fig. 2A**). As expected, acute AMPH administration induced significant increases in open-field locomotor activity (**Fig. 2B**). nucRNA-seq confirmed enrichment of PV+ and SST+ cell-type specific markers in the pulldowns relative to the UFs (**Fig. 2C)** and significant AMPH-dependent induction of the rapid PRG *Fos* in nuclei of PV+ neurons, SST+ neurons, and SPNs of the combined UF from both IPs (**Fig. 2D**).

**Figure 2:**
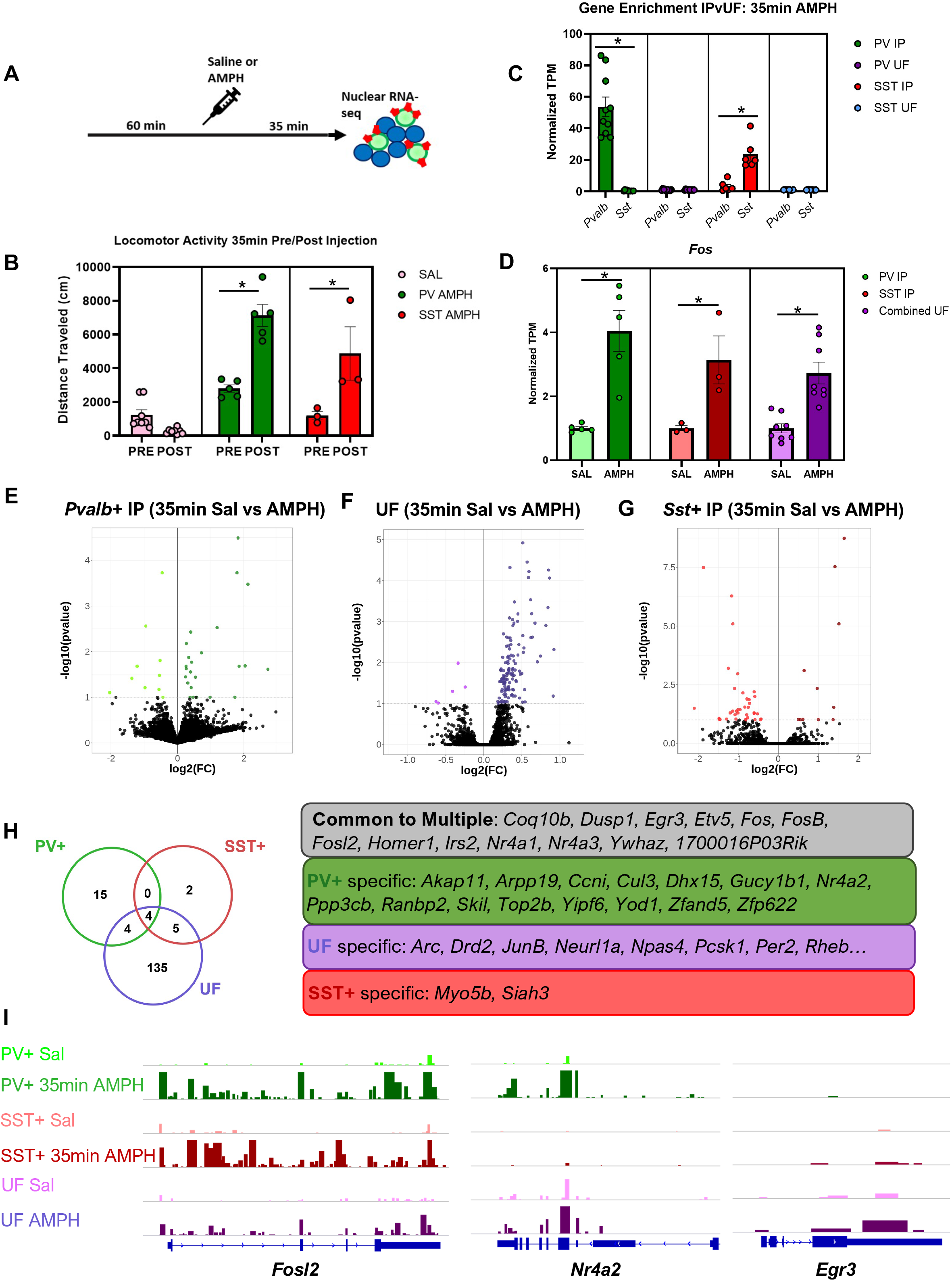
AMPH induces an overlapping program of rapid PRGs in distinct populations of NAc GABAergic neurons. **A)** Experimental timeline. Mice received saline (Sal) or amphetamine (AMPH, 3mg/kg, i.p.) after 60 min habituation to the open field. Brains were harvested 35 min later for nucRNA-seq. **B)** Summed locomotor activity in the open field 60 min before (Pre) and 30 min after (Post) i.p. injection of saline or 3mg/kg AMPH. *Pvalb-*Cre n=5/treatment; *Sst*-Cre n=3/treatment; combined UF n=8/treatment, Error bars indicate SEM. Two-way rmANOVA, *Pvalb-*Cre F (1, 11) = 93.18, p<0.0001, Bonferroni post-hoc AMPH Pre vs Post p<0.0001; *Sst-*Cre F (1, 9) = 24.91, p=0.0007), Bonferroni post-hoc AMPH Pre vs Post p=.0025. **C)** Validation of enrichment for cell-type specific marker transcripts (*Pvalb* or *Sst*) in nuclei recovered from each condition. TPM for each gene from Table S2 in the IP is shown normalized to UF TPM, Error bars indicate SEM. **D)** Quantification of NucRNA for the rapid PRG *Fos* in nuclei recovered from each condition from Table S2. *FDR<0.1, TPM normalized to SAL condition for each isolation, Error bars indicate SEM. **E-G**) Volcano plots of AMPH-regulated RNA at 30min post-injection in *Pvalb* IP (**E**, green), UF fractions (**F**, purple), or *Sst* IP (**G**, red). Black dots, AMPH vs SAL not significant; colored dots *FDR<0.1. Darker colors indicate genes induced by AMPH, lighter colors indicate genes repressed by AMPH. **H)** Venn diagram showing overlap of genes induced 35 min following AMPH in each population of nuclei. Common and cell-type specific induced genes are shown in the table at right. **I)** Representative NucRNA-seq tracks of PRGs *Fosl2*, *Nr4a2*, and *Egr3* in each population of nuclei 35 min after AMPH administration.

At this short time point after AMPH exposure, we detected a relatively small number of changes in gene expression in any of the cell types (**Fig. 2E-G, Table S2)**. In all three cell populations we observed an upregulation of a common set of rapid PRG TFs including members of the Fos and Nur families (**Fig. 2H**). These data are consistent with prior studies that found the overall rapid PRG transcriptional regulation program to be largely conserved between different types of neurons^25^. However, we did detect differential induction for specific family members in the rapid PRGs TF program, with *Nr4a2* showing induction in PV+ but not SST+ neurons (**Fig. 2I)**, and *Egr3* induction in the UF and SST+ neurons, but not PV+ neurons (**Fig. 2I)**.

Beyond the rapid PRG TFs, other genes rapidly induced by AMPH were largely divergent between the NAc GABAergic cell types we profiled, though they include gene products with known functions in plasticity (**Fig. 2H, Table S2**). Only in the UF fraction did we see induction of the canonical neuronal activity-regulated genes *Arc, Pcsk1, Per2,* and *Rheb*, many of which have been shown to function in SPNs to regulate cellular and behavioral responses to drugs of abuse^26, 27^. Only SST+ neurons showed AMPH-dependent induction of *Myo5b*, a calcium-regulated myosin motor that mediates recycling endosome trafficking in the context of LTP^28^. In PV+ neurons, many of the AMPH-induced genes are components of intracellular signaling pathways, though few of these genes have been studied as stimulus-regulated in the past. However, several have been implicated in neurological or psychiatric disorders, including the ubiquitin ligase *Cul3* in ASD and schizophrenia^29^, the nucleocytoplasmic transport protein *Ranbp2* in neurodegeneration^30^, and the topoisomerase *Top2b* for long gene regulation in ASD^31^. These data confirm that we can discover novel molecular programs of neuronal plasticity by comparing AMPH-regulated genes among distinct GABAergic cell types in the NAc.

### AMPH induces cell-type specific late gene programs in NAc interneurons

To identify the delayed PRGs and SRGs regulated by AMPH, we used INTACT to purify PV+ or SST+ interneurons from the NAc of mice 3hrs following an injection of either saline or AMPH in an open field chamber (**Fig. 3A**). We saw a robust effect of AMPH administration on locomotor activity in the open field (**Fig. 3B**), and we confirmed by nucRNA-seq that the interneuron markers *Pvalb* and *Sst* were enriched in their respective IP fractions when compared to the combined UF and to each other (**Fig. 3C**). We verified AMPH-dependent induction in the UF of *Bdnf*, which is an established delayed PRG^24^ and we confirmed the absence of *Bdnf* signal in the PV+ and SST+ nuclei harvested after AMPH exposure, as *Bdnf* is not inducible in GABAergic interneurons^32^ (**Fig. 3D**).

**Figure 3:**
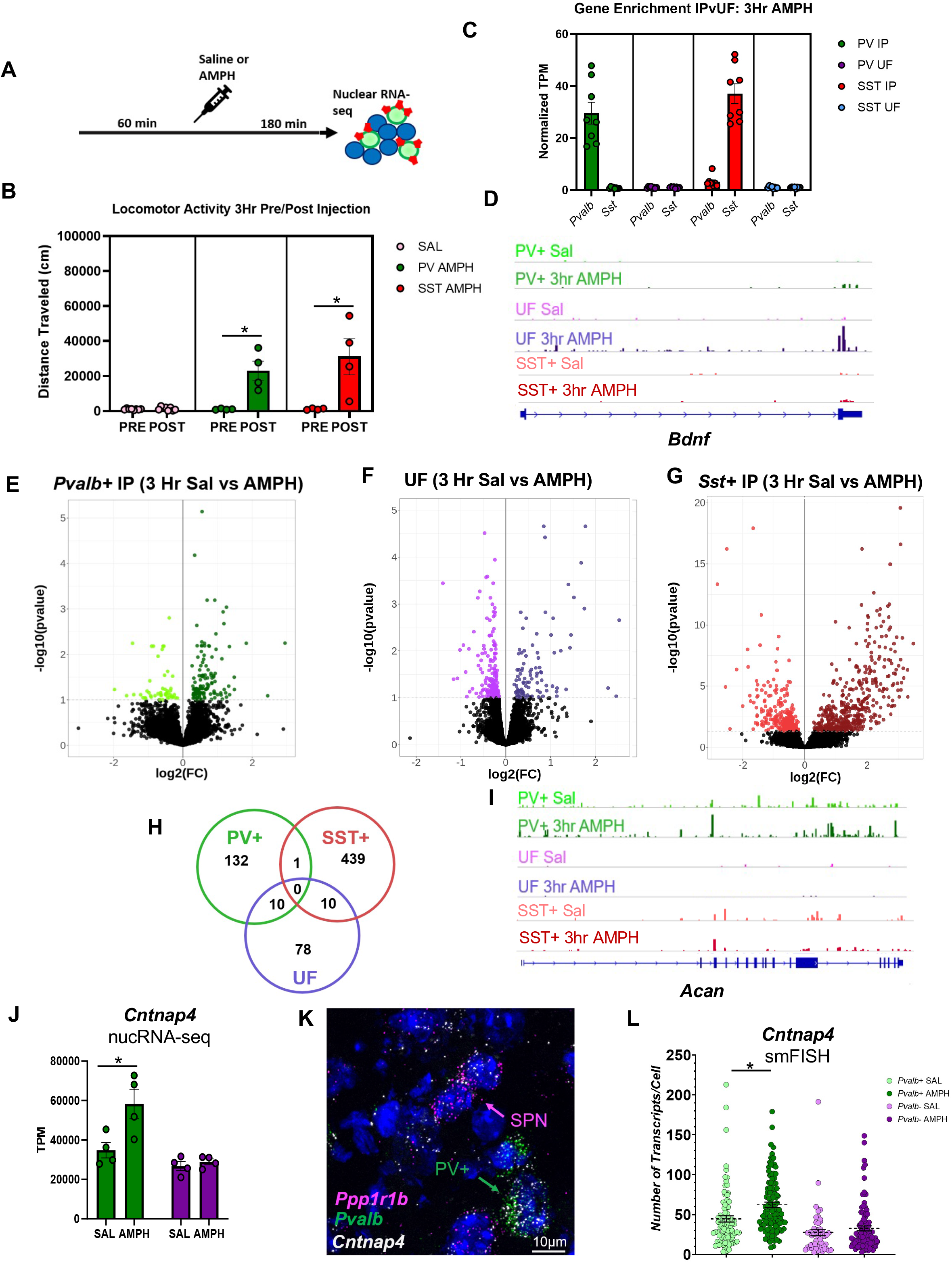
Cell-type specific programs of gene expression regulated 3 hrs after AMPH administration in different populations of NAc GABAergic neurons. **A)** Experimental timeline. Mice received an injection of saline (Sal) or amphetamine (AMPH, 3mg/kg, i.p.) after 60 min habituation in the open field. Brains were harvested 180 min (3 hr) later for nucRNA-seq. **B)** Locomotor activity 60 min before (pre) and 180 min after (post) i.p. injection of saline (light green) or 3mg/kg AMPH (dark green). *Pvalb-*Cre n=4/treatment condition; *Sst*-Cre n=4/treatment condition; Combined UF n=8/treatment condition; Two-way rmANOVA, *Pvalb-*Cre F (1, 10) = 33.43, p=0.0002, Bonferroni post-hoc AMPH Pre vs Post p<0.0001; *Sst-*Cre F (1, 10) = 17.77, p=0.0018), Bonferroni post-hoc AMPH Pre vs Post p=0.0008, Error bars indicate SEM. **C)** Validation of enrichment for cell-type specific marker transcript expression in nuclei recovered from each condition. TPM normalized to UF for each isolation. *Pvalb-*Cre IP n=4/condition; *Sst*-Cre IP n=4/condition; Combined UF n=8/condition, Error bars indicate SEM. **D)** Example Tracks of RNA for the delayed primary-response gene *Bdnf* in nuclei from each of the conditions. TPM normalized to SAL condition for each isolation. *Pvalb-*Cre IP n=4/treatment condition; *Sst*-Cre IP n=4/treatment condition; Combined UF n=8/treatment condition; *FDR<0.1. Y-axis proportionally adjusted for differential depth in SST Sal/AMPH resulting from PE sequencing. **E-G**) Volcano plots of AMPH-regulated gene expression at 180 min post-injection in *Pvalb*-Cre IP (E, green) *Sst*-Cre IP (G, red) or the combined UF fractions (F, purple). Black dots, SAL vs AMPH not significant; colored dots, using DESeq2 FDR<0.1. Darker colors indicate genes induced by AMPH; lighter colors indicate genes repressed by AMPH at 3 Hr post-administration. *Pvalb-*Cre IP n=4/treatment condition; *Sst*-Cre IP n=4/treatment condition; Combined UF n=8/treatment condition; *FDR<0.1. **H)** Venn diagram showing overlap of genes induced 3 Hr following AMPH in each population of nuclei. **I)** Representative nucRNA-seq track for example cell-type specific AMPH-induced gene *Acan*. **J-L)** Cell-type specific induction of *Cntnap4* by AMPH in *Pvalb*+ neurons of the NAc. J) nucRNA-seq quantification of *Cntnap4* TPM in *Pvalb-*Cre IP n=4/treatment condition; *Pvalb-*Cre UF n=4/treatment condition; *FDR<0.1 for +/− AMPH treatment. K) smFISH for *Cntnap4* and *Pvalb* in NAc. Scale bar = 10µm. L) Quantification of *Cntnap4* smFISH in *Pvalb*+ and *Pvalb*-nuclei; Two-way ANOVA, F (1, 332) = 9.093, p=0.0028, Bonferroni post-hoc *Pvalb*+ Sal vs AMPH p=.0017.

We identified 143 AMPH-induced genes in NAc PV+ interneurons, 450 in SST+ interneurons, and 98 in the combined UF (**Fig. 3E-G; Table S3**). In contrast to the overlapping programs of rapid PRGs induced by AMPH across cell types, the delayed gene programs were almost completely distinct (**Fig. 3H**). Gene Ontology (GO) analysis of the AMPH-regulated genes showed gene categories related to multiple signal transduction pathways in all three cell populations, suggesting, as we expected, that all these cells were experiencing intracellular adaptations to acute pharmacological stimulation (**Fig. S2A-F**). However, we were particularly interested to see upregulation selectively in PV+ neurons of genes in categories that affect synapse structure and function (positive regulation of synaptic transmission, positive regulation of synapse assembly, chemical synaptic transmission) and excitability (potassium ion transport, positive regulation of cytosolic calcium ion concentration) (**Fig. S2A**). By contrast, the downregulated pathways in PV+ neurons were predominantly related to general metabolic and biosynthetic pathways **(Fig. S2D)**. These data suggest that following AMPH exposure, PV+ neurons may divert basal resources to functionally remodel their connectivity and inhibitory efficacy within local circuits, which could change the impact of their activation.

The category of cell adhesion was significantly enriched in both PV+ and SST+ AMPH-induced genes, including some genes already known to influence properties of interneuron synapses. For example, *Acan* encodes the perineuronal net (PNN) protein Aggrecan (**Fig. 3I).** Knockout of Aggrecan disrupts PNNs and switches PV+ neurons to a high plasticity state *in vivo*^33^, and some prior studies have suggested roles for PNNs in neural plasticity induced by drugs of abuse^34^. *Cntnap4* (**Fig. 3J-L)** is a member of the neurexin superfamily of cell adhesion proteins that promotes presynaptic release of GABA from PV+ interneurons. Knockout of *Cntnap4* augments dopamine release in the NAc and dampens inhibition from PV+ interneurons^35^.

Notably, although the AMPH-dependent induction of genes is largely cell-type specific, we observed that most of the inducible genes were expressed under basal conditions in more than one NAc neuronal population. For example, we detected significant enrichment of *Acan* in both PV+ and SST+ interneurons compared with the UF but find a selective induction of *Acan* by AMPH only in PV+ neurons and not SST+ neurons (**Fig. 3I**). For *Cntnap4,* by RNAseq we observed expression in both PV+ and SST+ neurons as well as the UF but only saw significant AMPH-induced increases in *Cntnap4* in PV+ neurons (**Fig. 3J)**. We validated this observation with quantitative FISH, confirming that both *Ppp1r1b*+ SPNs and *Pvalb*+ interneurons within the NAc express *Cntnap4* **(Fig. 3K)** but only *Pvalb*+ interneurons show a significant increase in *Cntnap4* signal following AMPH when compared to surrounding PV-, *Cntnap4*-expressing cells. (**Fig. 3L**).

### Stable chromatin accessibility landscapes in NAc cells following AMPH exposure

Because enhancer usage can be highly cell-type specific even for genes with broad expression patterns^36^, we examined chromatin accessibility in NAc interneurons to determine possible transcriptional mechanisms of cell-type specific AMPH regulation. To characterize chromatin accessibility for TF binding genome-wide, we used the Assay for Transposase-Accessible Chromatin (ATAC-seq) on neuronal nuclei purified by INTACT. In AMPH-naïve mice, PV+ and SST+ interneurons have a unique and replicable chromatin accessibility landscape that distinguishes them from each other and from the GABAergic SPNs that predominate in the UFs **(Fig. 4A-C)**. When compared to their respective UFs, we find >60,000 differentially accessible regions of chromatin uniquely accessible in PV+ and SST+ interneurons genome wide. (**Table S4).** Conversely, we find 46,348 regions that are uniquely accessible in the combined, SPN-enriched UF. More modestly, we find ∼5000 unique differentially accessible regions between immunoprecipitated PV+ and SST+ interneurons. Consistent with prior studies^37, 38^, only a small fraction (<10%) of the differentially accessible sites were found at gene promoters, whereas the vast majority occur at inter- and intragenic sites that are likely to function as distal transcriptional regulators (**Fig. 4D**).

**Figure 4:**
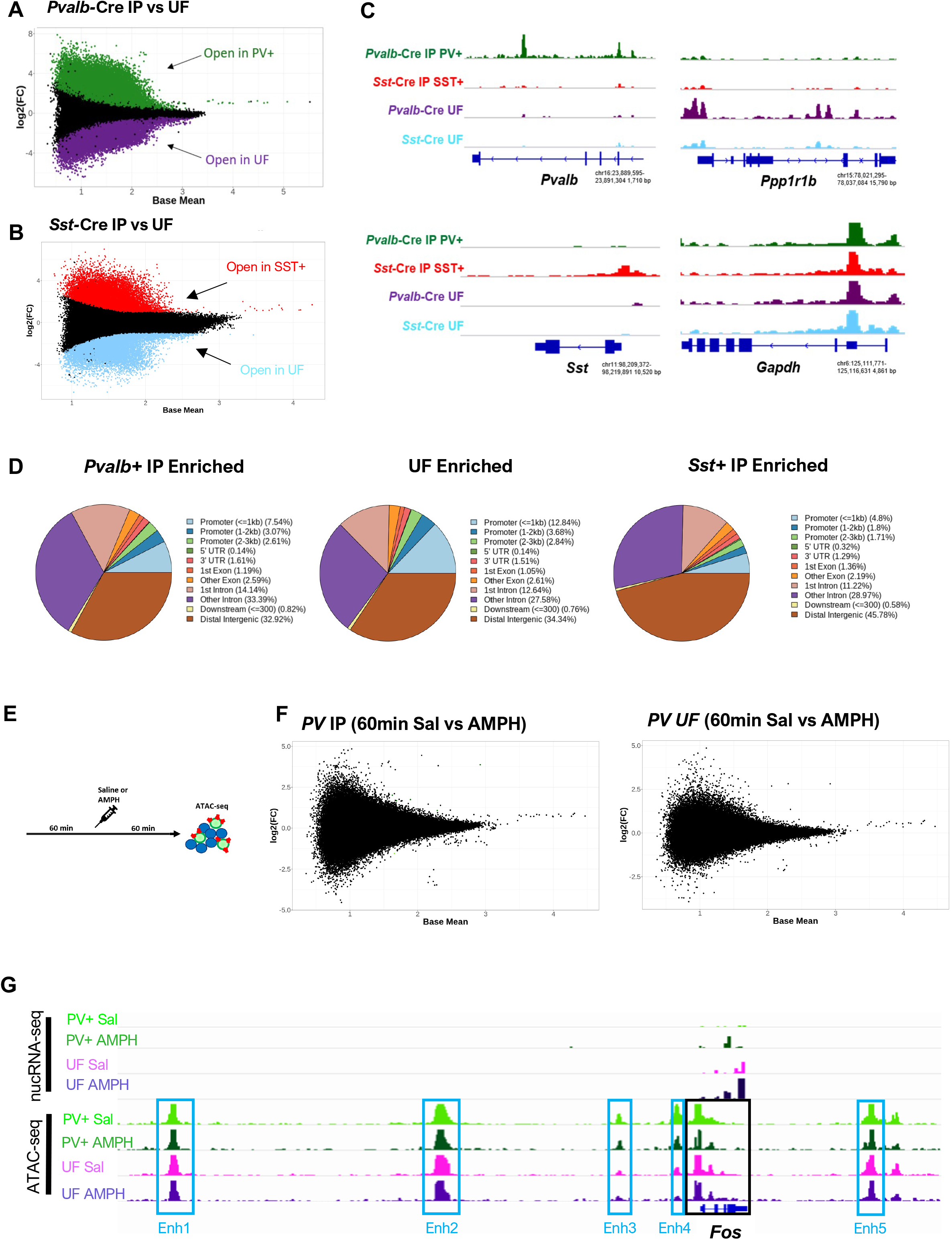
Cell-type specific and post-AMPH chromatin accessibility in NAc neurons. **A-B)** MA plots of cell-type specific differential chromatin accessibility in each population of isolated nuclei using DeSeq2 *FDR<0.05. *Pvalb-*Cre IP vs *Pvalb-*Cre UF, Green points indicate regions significantly differentially accessible in PV+ nuclei; Purple points indicate regions significantly differentially accessible in *Pvalb*-Cre UF nuclei (A); *Sst*-Cre IP, red *Sst-*Cre IP vs *Sst-*Cre UF, blue points indicate regions significantly differentially accessible in SST+ nuclei; Light blue points indicate regions significantly differentially accessible in *Sst*-Cre UF nuclei (B). **C)** Example tracks of cell-type specific accessible regions in the vicinity of cell-type marker genes in each isolated cell fraction. *Gapdh* track included as a commonly accessible reference gene in all cell types. Y-axis is consistent across all tracks for each gene. **D)** Pie chart depicting relative genomic location (Promoter, Gene body/Intragenic, Downstream, or Distal Intergenic) of cell type specific (*Pvalb-*Cre IP, *Sst*-Cre IP, Combined UF) differentially accessible chromatin regions enriched in each fraction. **E)** Experimental timeline. Mice received an injection of saline (Sal) or amphetamine (AMPH, 3mg/kg, i.p.) after 60 min habituation in the open field. Brains were harvested 60 min later for ATAC-seq**. F)** MA plots of AMPH-induced differential chromatin accessibility in each population of isolated nuclei 60 min post administration using DeSeq2 *FDR<0.05; *Pvalb-*Cre IP n=4/treatment conditions *Pvalb-*Cre UF n=4/treatment condition. **G)** Representative genomic tracks of nuc-RNAseq and ATAC-seq from Sal control and AMPH-treated samples on and in the vicinity of the *Fos* gene. nucRNA-seq tracks show AMPH-induced expression of *Fos* at 35min in both PV+ and UF cell populations. ATAC-seq depicts chromatin accessibility at the *Fos* gene and at its five known enhancer regions outlined in blue.

Some studies have reported dynamic changes in chromatin accessibility following stimuli that lead to the activation of rapid PRG TFs^39–40^. These changes may reflect the concerted eviction of histones by RNA polymerase II (RNAPolII) during active transcription or the recruitment of rapid PRG TFs to regulatory elements driving subsequent chromatin remodeling. Given that we observed robust and overlapping programs of rapid PRG TF induction in all our NAc nuclear fractions following AMPH (**Fig. 2**), we asked whether this induction was associated with changes in chromatin accessibility in PV+ neurons or the corresponding SPN-enriched UF.

We administered either saline or AMPH to mice in the open field and harvested PV+ neurons by INTACT 60min later (**Fig. S3A**). The accessibility landscape of PV+ interneurons in both conditions was comparable to that of drug-naïve mice and clearly distinguished from accessibility in the UF **(Fig. S3B)**. However, acute AMPH exposure did not induce any substantial changes in chromatin accessibility either in PV+ interneurons or in the UF **(Fig. 4E-F; Table S5)**. This stability of chromatin accessibility was evident even at known regulatory elements controlling production of the rapid PRG TFs despite their AMPH-induced transcription, as shown for *Fos* (**Fig. 4G**) and other rapid PRGs (**Fig S3C)**. To determine whether changes in chromatin accessibility might require repeated exposure to AMPH, we next performed ATAC-seq following a 7d repeated AMPH locomotor sensitization paradigm (**Fig. S3D**). 24hrs following the final AMPH administration we harvested PV+ nuclei by INTACT for ATAC-seq. This stimulus was associated with significant differential expression of 361 transcripts within PV+ cells (**Fig. S3E, Table S6)** across various GO categories (**Fig. S3F)**. 6 of the chronic AMPH-regulated genes overlapped the set changed 3hr after acute AMPH, suggesting persistent changes in transcription following repeated AMPH-exposure **(Table S6)**. Nonetheless, we again observed no substantial changes in accessibility in either the PV+ or the UF fractions after repeated AMPH exposure **(Fig. S3G; Table S7)**.

### Single-nucleus RNA-Seq of PV+ nuclei reveals rapid PRG induction in multiple PV+ subtypes

By immunostaining we found that only a small percentage of PV+ neurons (∼15%) show robust, AMPH-dependent increases in Fos protein levels (**Fig. S4A,B**). We thus considered the possibility that heterogeneity in our purified PV+ nuclear fraction could mask chromatin accessibility dynamics in a subset. To determine whether molecularly identifiable PV+ neuron subtypes were distinguishable within our purified population, we used fluorescence-activated nuclear sorting (FANS) to isolate Sun1-GFP tagged nuclei from *Pvalb*-Cre mice for single nuclear RNA sequencing (snRNA-seq) on the 10X Genomics platform. We harvested and pooled NAc nuclei from mice (Sal n=3, AMPH n=4) 35min following an injection of either Sal or AMPH in the open field **(Fig. S4C,D)**. Prior to FANS, we incubated nuclei from each condition with unique lipid-modified oligonucleotides (LMOs)^42^ to allow for multiplexing and *post-hoc* bioinformatic identification of nuclei from the saline and AMPH-treated mice.

After filtering GFP captured cells for *Pvalb* expression we successfully recovered a total of 787 PV/GFP+ nuclei with a mean read depth of 6,930 counts per nucleus and a median 2,968 genes sequenced per nucleus **(Fig. S4E; Table S8)**. We performed dimensionality reduction via principal components analysis (PCA) for generation of a UMAP that defined 7 clusters of PV+ neurons (**Fig. S4F-H**). These clusters all expressed the GABA synthesizing enzyme *Gad1*, which is enriched in PV+ interneurons^43^ (**Fig. S4I**). None of the clusters contained the glial markers *Gfap* and *Sox10*, or the SPN marker *Ppp1r1b,* indicating that we had little contamination from other major cell types of the NAc in our filtered population.

The top two genes contributing the greatest amount of cell-to-cell variance across the *Pvalb*-expressing clusters were Adenosine Deaminase RNA Specific B2 (*Adarb2*) and cell surface heparan sulfate proteoglycan Glypican-5 (*Gpc5*) **(Fig. 5C),** both of which were most highly expressed in cluster 4. It was surprising to find *Adarb2* coexpressed in *Pvalb*+ neurons, because at least for cortical interneurons, *Adarb2* has been characterized as a marker of interneurons that originate from the caudal ganglionic eminence during development, whereas *Pvalb*+ neurons are thought to originate from the medial ganglion eminence^44^. We used FISH on coronal sections of NAc from mouse brain to confirm coexpression of *Adarb2* in a subset of *Pvalb*+ interneurons **(Fig. 5D),** whereas no colocalization was observed in cortex from the same mice **(Fig. S4J).** We subset our nuclei into *Pvalb*+ and *Adarb2*+/− identities and performed differential expression analysis using Wilcoxon rank-sum tests to identify transcripts significantly differentially expressed between these two predefined clusters (**Fig. S4K**).

**Figure 5:**
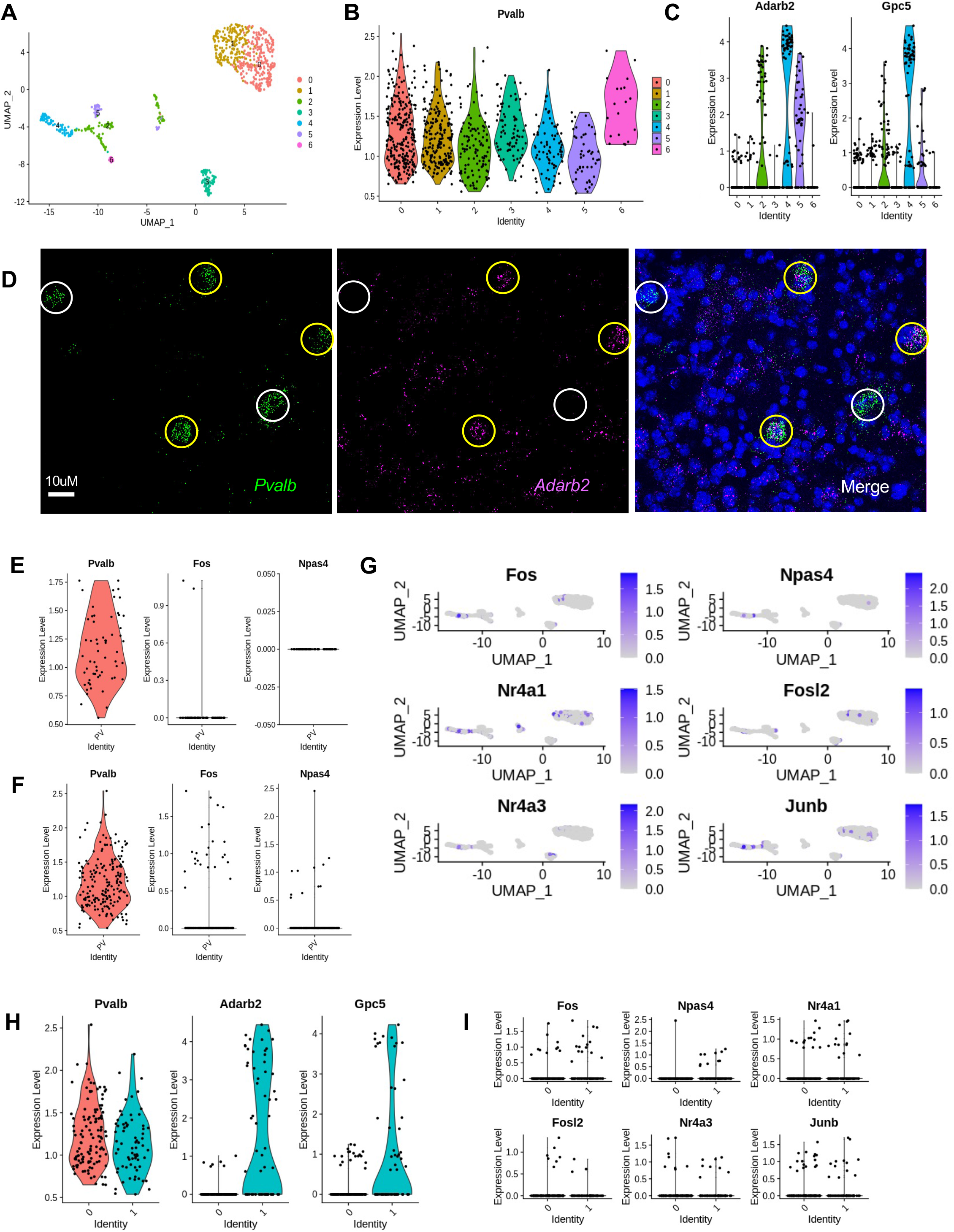
Single nucleus RNA-seq reveals molecular heterogeneity of PV+ interneurons in the NAc. **A)** Seven-cluster UMAP projection of snRNAseq data from *Pvalb*-Cre nuclei isolated with FANS; n= 787 nuclei after normalization, scaling, and filtration based on detectable polyadenylated *Pvalb* transcripts. **B)** Violin plot of *Pvalb* log-normalized expression levels in nuclei across the 7 UMAP projection clusters. **C)** Violin plots of log-normalized expression levels of *Adarb2* and *Gpc5* in nuclei across the 7 UMAP projection clusters. **D)** Fluorescent in situ hybridization using probes against *Pvalb* and *Adarb2* in the mouse NAc exhibiting partial colocalization of *Pvalb* and *Adarb2* RNA in single cells; Yellow circles indicate *Pvalb*/*Adarb2* co-positive cells, white circles indicate *Pvalb+/Adarb2-*cells. **E)** Violin plots of log-normalized expression levels of *Pvalb*, *Fos*, and *Npas4* in nuclei confirmed positive for Multi-seq LMO 5 (top, SAL treated n=60) or LMO6 (bottom, AMPH-treated, n=187). **G)** Feature plots depicting nuclei with detectable transcripts of various PRGs across the 7 UMAP projection clusters. **H)** Violin plot of log-normalized expression levels of *Pvalb*, *Adarb2*, and *Gpc5* in nuclei confirmed positive for Multi-seq LMO 6 (AMPH) in two-cluster UMAP projection of snRNA-seq data from *Pvalb*-Cre nuclei. 0, *Adarb2-/Gpc5-*; 1, *Adarb2+/Gpc5+*. **I)** Violin plot of log-normalized expression levels of various PRGs in two-cluster UMAP projection of snRNA-seq data from *Pvalb*-Cre nuclei confirmed positive for Multi-seq LMO 6 (AMPH) clustered as in **H**.

To determine whether AMPH-dependent gene induction was occurring within specific subsets of PV+ neurons, we first deconvolved the multi-seq tags to confirm that we could detect induction of rapid PRG TFs in nuclei from the brains of AMPH-treated mice in our snRNA-seq data. The LMO barcodes were successfully amplified in a subset of our sequenced libraries allowing us to confirm enrichment of rapid PRGs in nuclei of mice exposed AMPH relative to those exposed to saline (n=60 Sal, n=187 AMPH) **(Fig. 5E,F; Fig. S4L,M)**. We observed expression of rapid PRGs in all 7 PV+ clusters, suggesting that the response to AMPH was not limited to a single cluster **(Fig. 5G)**. We then created identities to subset our *Pvalb+* nuclei into *Adarb2*+/− groups (**Fig. 5H**), however, expression of rapid PRGs was similar in both subsets (**Fig. 5I)**. Taken together these data confirm that a fraction of PV+ neurons in the NAc respond to AMPH with a rapid PRG transcriptional response. However, this fraction is not a molecularly defined subset of PV+ interneurons, suggesting it is more likely to be a result of differential functional or developmental connectivity.

### Cell-type specific transcriptional regulation of AMPH-dependent genes

Although we saw no AMPH-dependent changes in chromatin accessibility, we did observe cell-type specific regions of accessible chromatin between our isolated cell types that correlated with cell-type specific AMPH-dependent transcriptional regulation **(Fig. 6A)**. As such, we next asked if the unique landscapes of cell-type specific chromatin accessibility in GABAergic NAc neuronal types could reveal differential binding sites for transcription factors (TFs) that mediate cell-type specific transcriptional responses to acute AMPH.

**Figure 6:**
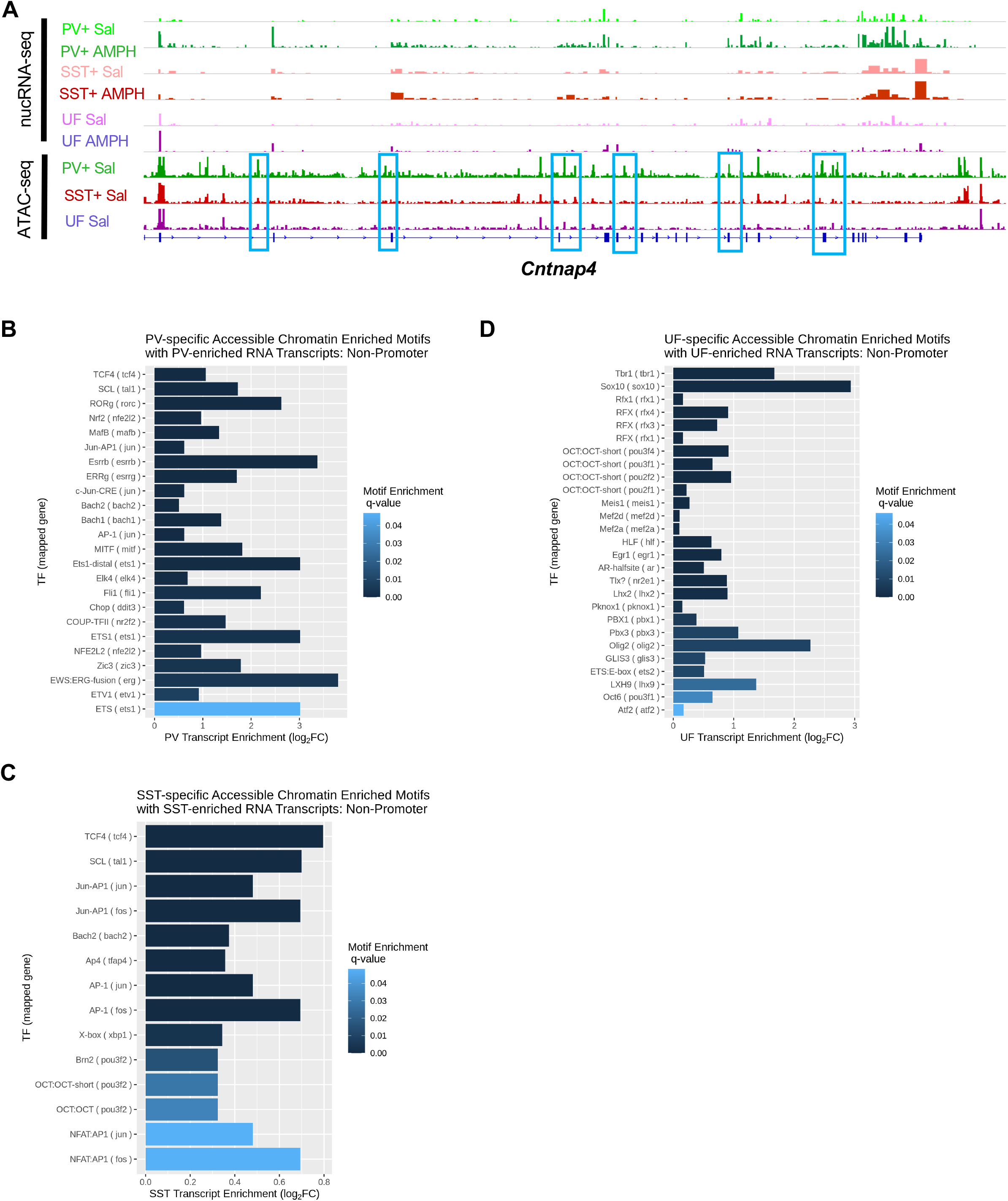
Motif analysis of differentially accessible chromatin near AMPH-regulated genes suggests transcriptional mechanisms of gene regulation in NAc neurons. **A)** Representative genomic tracks of nuc-RNAseq and ATAC-seq from Sal control only (ATAC) or Sal and AMPH-treated samples (nuc-RNAseq) in the vicinity of the *Cntnap4* gene. nucRNA-seq tracks show significant AMPH-induction of *Cntnap4* RNA specific to the PV+ cell population. ATAC-seq depicts regions within the *Cntnap4* gene significantly differentially accessible within PV+ interneurons (*Pvalb-*Cre IP) outlined in blue. **B-D)** Enriched Transcription Factor Motifs as determined by HOMER at cell-type-unique differentially accessible inter- (+/− 50Kb) and intragenic chromatin regions at genes induced by AMPH at 3Hrs in each cell fraction, *q<0.05; Enriched motifs are plotted against log2FC enrichment at baseline of cognate RNA transcript in each isolated cell type or UF; *Pvalb*-Cre IP vs UF (**B**), *Sst-*Cre IP vs UF (**C),** Combined UF vs Combined IP (*Pvalb*-Cre+*Sst-*Cre IP)(**D**).

For all the genes induced in PV+ interneurons, SST+ interneurons or the combined UF at 3hr post AMPH, we identified regions of differential chromatin accessibility between cell types at promoters (defined as 1kb on either side of the transcription start site) and putative enhancers (including the gene body and distal regions +/− 50kb on either side of the gene but excluding the promoter). We searched these regions for enriched transcription factor binding motifs, matched the motifs to TF families (**Fig. S5A)** and then identified those TFs that showed cell-type enriched **(Fig. 6B-D)** or AMPH-regulated expression **(Fig. S5B-D)** in the cell type that displayed open chromatin relative to other NAc cell types.

We found more diversity of enriched TF binding sites in the enhancers compared with the promoter regions, consistent with prior evidence that intragenic and distal enhancers are major regulators of cell-type specific gene expression^5^. Thus, we focused our analysis to identify TFs that bind these putative enhancers. In all three cell types we identified enrichment of binding sites for multiple TFs that show cell-type specific expression (**Table S9**). TCF4 was a top hit in both PV+ and SST+ neurons (**Fig. 6B,C)**. Although TCF4 is broadly expressed in the cortex, within striatal regions its expression is limited to GABAergic interneurons^45^. TCF4 interacts with β-catenin to regulate gene expression downstream of the Wnt signaling pathway. Little is known about the functions of Wnt signaling in addiction, but at least one study has found increased nuclear levels of β-catenin in the NAc following cocaine^46^ and another showed important functions for NAc β-catenin in resilience to chronic stress^47^. PV+-specific enhancers also show enrichment for ETV1, a TF that controls intrinsic firing properties of PV+ cortical interneurons, where its abundance is activity-regulated^48^. By contrast, differentially accessible enhancers in the SPN-enriched UF fraction showed enrichment of binding sites for members of the RFX and MEF2 families of TFs among others **(Fig. 6D**). *Rfx1,3,* and *4* are enriched in the UF fraction relative to the interneuron populations (**Fig. 6D; Table S1**) and *Rfx4* is rapidly induced by AMPH in the UF (**Fig. S5D; Table S2**). The psychostimulant-dependent regulation and function of the MEF2s in mediating cocaine-induced synapse plasticity in SPNs has been well described^49^.

These data suggest that the differential pattern of accessible enhancers near AMPH-regulated genes is maintained by cell-type specific control of the expression of TFs some of which are targets of regulation by psychostimulant-induced signaling cascades. However, we also found enrichment of binding sites in for rapid PRG TFs of the Fos, Jun, and Egr families (**Fig. S5B-D**), suggesting, as has been previously proposed, that these ubiquitous TFs work together with cell-type specific TFs to amplify programs of stimulus-regulated gene transcription^50^. The rapid induction of these TFs may drive the later cell-type specific programs of gene transcription by acting at cell-type specific sites of accessibility.

### Dysregulation of PV+ neuron gene regulation in MeCP2 Ser421Ala knockin mice

To begin to determine which genes in PV+ interneurons might contribute to behavioral responses to drugs of abuse, we assessed PV+ neuron gene expression in the NAc of MeCP2 Ser421Ala KI mice. These mice show both behavioral hypersensitivity to psychostimulants and altered AMPH-regulated Fos expression in NAc PV+ interneurons^18^, thus we hypothesized that gene expression differences in PV+ interneurons of these mice could reveal genes important for addictive-like behaviors. We used the RiboTag method^51^ to enrich for actively translating mRNAs from NAc PV+ neurons of MeCP2 WT and Ser421Ala KI mice **(Fig. 7A)**. We confirmed co-expression of the HA tagged ribosomal subunit with PV in the NAc (**Fig. 7B**) and we found enrichment of *Pvalb* mRNA in the immunoprecipitated fraction from both MeCP2 WT and Ser421Ala KI mice relative to each input (**Fig. 7C)**. Importantly, despite significant differences in the method of RNA isolation we saw substantial overlap in PV+-specific gene expression (**Fig. 7D**) comparing actively translating mRNAs isolated by ribosome pulldown (**Fig. 7A**) and the nascent RNAs isolated by nuclear pulldown (**Fig. 1A**).

**Figure 7:**
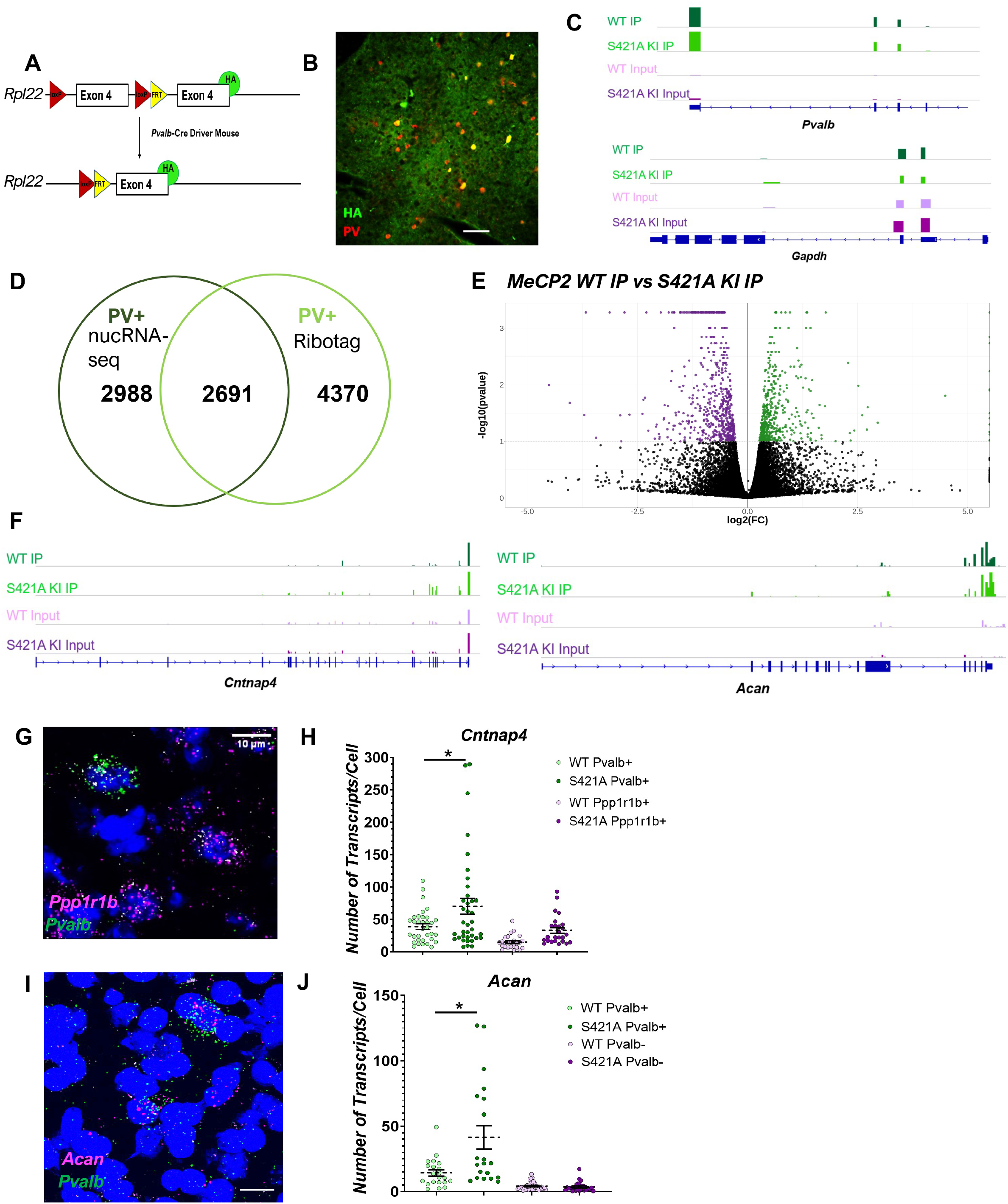
Gene dysregulation in NAc PV+ neurons of MeCP2 Ser421Ala knockin mice. **A)** Depiction of *Pvalb*-2A-Cre-dependent HA tagging to Rpl22 for cell-type specific (PV+) isolation of actively translating mRNA. **B)** IHC depicting specificity of HA tag expression with PV protein. **C)** Representative genomic tracks at the *Pvalb* gene of WT and S421 KI IP and Input fractions demonstrating significant enrichment of *Pvalb* RNA in the IP conditions. *Gapdh* gene track included as a commonly expressed reference gene in all fractions. Y-axis is consistent across all tracks for each gene. **D)** Venn diagram of overlapping, cell-type specific basal gene enrichment in PV+ cells as measured by nucRNA-seq (dark green) and Ribotag (light green) vs surrounding cells (*Pvalb-*Cre IP vs UF nucRNAseq, WT IP vs WT Input Ribotag). **E)** Volcano Plot of differentially dysregulated genes between MeCP2 WT vs KI immunoprecipitated fractions (IP) in naïve mice; Black dots, not significant; colored dots, FDR<0.1; n=15/genotype, pooled into 3 replicates of 5 mice each. **F)** Representative genomic tracks at the nucRNA-seq AMPH-induced genes *Acan* and *Cntnap4* of WT and Ser421Ala (S421A) KI IP and Input fractions. **G)** Representative image of FISH targeting *Pvalb*, *Ppp1r1b*, and *Cntnap4* mRNA. **H)** smFISH quantification of *Cntnap4* transcript number in *Pvalb+* or *Ppp1r1b+* nuclei (n=34 WT PV, 37 KI PV+ nuclei, 24 WT SPN nuclei, 24 KI SPN nuclei); Two-way ANOVA, F (1, 117) = 8.943, *p=0.0034, Bonferroni post-hoc *Pvalb* WT vs S421A p=.0226. **I)** Representative image of FISH targeting *Pvalb* and *Acan* mRNA. **J)** smFISH quantification of *Acan* transcript number in *Pvalb+* or *Ppp1r1b+* nuclei (n=21 WT *Pvalb*+, 20 KI *Pvalb+* nuclei, 37 WT *Ppp1r1b+* nuclei, 28 KI *Ppp1r1b+* nuclei) Two-way ANOVA, F (1, 102) = 13.47, *p=0.0004, Bonferroni post-hoc *Pvalb*+ WT vs S421A p<.0001.

We found 1082 transcripts differentially expressed in the PV+ interneurons of MeCP2 Ser421Ala KI mice compared with their WT littermates **(Fig. 7E; Fig. S6A; Table S10)**, a subset of which overlapped with our AMPH-induced PV+ program of delayed PRGs and SRGs. PV+ neurons of the MeCP2 Ser421Ala KI mice showed elevated expression of the PV-specific, AMPH-inducible genes *Cntnap4*, *Clstn2,* and *Acan* **(Fig. 7F-J; Table S10)**, whereas expression of the canonical housekeeping gene *Gapdh* did not differ by genotype **(Fig. S6B-C)**. Taken together, these data implicate these synaptic cell-adhesion gene products as promising candidates to modulate behaviorally-relevant properties of PV+ interneurons following exposure to psychostimulants.

## Discussion

In this study, we conducted cell-type specific RNA and chromatin sequencing to provide comprehensive identification of the *in vivo* gene regulatory responses induced by AMPH in NAc interneurons of adult mice. Our data show that transcriptional changes induced by AMPH are largely unique to specific GABAergic cell-types. We did not observe dynamic changes in chromatin accessibility following AMPH exposure, however we did find evidence that differential accessibility of transcriptional enhancers correlates with cell-type specific responses to AMPH. Finally, we identified a set of genes selectively regulated by AMPH in PV+ neurons that show altered expression in a mutant mouse strain that displays enhanced behavioral sensitivity to psychostimulants, suggesting potential for the functional importance of these gene expression programs for the expression of addictive-like behaviors.

Artificially enhancing or depressing the function of NAc PV+ interneurons modulates the expression of addictive-like behaviors^13, 14^. However, whether these neurons undergo transcriptional plasticity in response to psychostimulant exposure was unknown. Our data now identify hundreds of genes in NAc PV+ interneurons that show significant changes in their expression following acute or repeated AMPH exposure. Even though we were comparing gene regulatory programs between multiple GABAergic cell types within the NAc of individual mice, the genes regulated in each kind of neuron were largely distinct. These data suggest that even similar kinds of neurons experience distinct forms of cellular plasticity in response to a common stimulus, extending results of previous studies comparing more diverse cell types^25, 32^,

Examination of the PV+ specific AMPH-regulated gene expression program shows induction of cell adhesion proteins that localize to both pre- and postsynaptic sites. Taken together with the evidence that PV+ neurotransmitter release is positively correlated with the expression of addictive-like behaviors^13, 14^, these data suggest that PV+ interneurons may enhance their connectivity within NAc circuits following psychostimulant exposure. We validated PV+ interneuron specific induction of *Cntnap4*, a member of the neurexin superfamily that functions in presynaptic PV+ interneuron terminals to promote inhibitory synaptic strength by limiting the size of the synaptic cleft^35^. We observed PV+ specific induction of *Acan*, encoding aggrecan, a PNN component that plays an important role in organizing postsynaptic protein complexes in PV+ neurons^33, 52^. In addition, we see AMPH-induced PV+ neuron upregulation of *Clstn2*, calsyntenin-2, a member of the cadherin family of cell-adhesion molecules that functions to increase inhibitory synapse number through a mechanism that may involve interactions with the neurexin family of synapse organizing proteins^53, 54^. Expression of all three of these genes is elevated in PV+ neurons of AMPH-naïve MeCP2 Ser421Ala KI mice compared with their WT littermates, providing a potential mechanism for the enhanced behavioral sensitivity of these mice to psychostimulant drugs of abuse. Cell-type specific conditional knockouts of these gene products will be informative for their functional importance in addictive-like behaviors.

Cell adhesion genes as a category were also regulated in NAc SST+ interneurons following acute AMPH exposure, though the specific genes that were targets of regulation in the two cell types did not overlap. A prior study using FANS to isolate SST+ nuclei from the NAc also observed regulation of the cell adhesion genes *Ank3* and *Nrcam* after chronic cocaine exposure^8^. Like PV+ neurons, NAc SST+ interneuron activity is positively associated with the expression of locomotor sensitization and conditioned place preference after cocaine^8^, suggesting that enhancing local inhibition in the NAc could be a common circuit mechanism of addictive-like behaviors even if the molecular mediators of that state differ by interneuron cell type.

Although we observed largely cell-type specific programs of gene regulation by AMPH, many of the genes whose expression changed following AMPH exposure were expressed under control conditions in more than one NAc cell type. For example, we validated by smFISH that *Cntnap4* is expressed in both *Ppp1r1b*+ SPNs and *Pvalb*+ interneurons of the NAc but only induced by AMPH in the *Pvalb*+ population. We also find genes such as *Acan* that are expressed in both PV+ and SST+ neurons, but only AMPH-induced in PV+ neurons. Only one prior study directly compared differential stimulus-regulation of gene expression between populations of interneurons in a single brain region, and they limited their analysis to only those gene products that were only expressed at baseline in a single class of interneurons relative to other cells in the population^55^. Thus, much remains to be learned about the underlying mechanisms that confer cell type specificity on stimulus-dependent regulation.

Our chromatin data suggest that the differential accessibility of intra- and intergenic enhancers underlies the differences in the AMPH responsivity of genes between cell types. Although recruitment of the RNA polymerase to the proximal promoter region of a gene is ultimately required for the activation of transcription, it is distal enhancers that mediate celltype-, developmental stage-, and stimulus-specific modulation of transcription^5^. The link between enhancer activity and chromatin accessibility reflects the differential binding of TFs at these regulatory elements. Indeed, when we examined the DNA sequences of putative enhancers near our cell-type specific AMPH-regulated genes, we observed enrichment of binding sites for numerous TFs that display cell-type specific patterns of expression. In this manner, the pattern of available enhancers would be permissive for the ability of a gene to show stimulus-dependent regulation. However, these differentially accessible regions were also enriched for binding sites for rapid PRG TFs, suggesting that the common induction of this rapid program in all AMPH responsive cells could instruct differential programs of stimulus responsive transcription by collaborating with cell-type specific TFs, similar to the mechanisms proposed for neuronal activity-dependent regulation of development^56^.

Is there a role for chromatin plasticity in AMPH-dependent regulation of interneuron gene expression? Some studies have shown intriguing evidence that activity-dependent induction of rapid PRG TFs can drive the formation of new regions of accessible chromatin, leaving a lasting mark on the chromatin landscape that could potentially function as a form of epigenetic memory^39–40^. We did not find significant changes in chromatin accessibility in PV+ interneuron nuclei or the SPN-enriched nuclei of the UF either following acute or repeated AMPH, though we cannot rule out that these changes could have occurred in a small subset of neurons. However, accessibility is only one measure of chromatin state. Previously we have shown that Ser421 phosphorylation of the methyl-DNA binding protein MeCP2 is selectively induced in NAc PV+ neurons following AMPH exposure^16^, and here we have identified a program of gene expression that is dysregulated in NAc PV+ neurons of mice bearing a non-phosphorylatable Ser421Ala mutation knocked into the *Mecp2* gene. MeCP2 is highly abundant in neurons and it binds globally across CpG and CpA methylated regions of the genome, yet acts locally at transcription start sites to control transcriptional initiation^57, 58^. Null mutations in MeCP2 are associated with cell-type specific changes in heterochromatin compaction and the sub-nuclear distribution of certain post-translationally modified histones^59^. Although the consequences of MeCP2 Ser421 phosphorylation on nuclear architecture is unknown, the ability of MeCP2 to nucleate protein complexes that mediate gene repression has been shown to be modulated by phosphorylation at Thr308, impeding the interaction of MeCP2 with the NCoR repressor complex^60^. Future studies using measures of chromatin architecture that can be scaled to the single cell level^61, 62^ offer a promising approach to discovering novel mechanisms of AMPH-dependent chromatin regulation in NAc interneurons.

## Methods

### Animals

We performed all procedures under an approved protocol from the Duke University Institutional Animal Care and Use Committee. We used the following mouse strains: *Pvalb*-IRES-Cre (B6.129P2-*Pvalb*^tm1(cre)Arbr/^J, RRID:IMSR JAX:017320); *Sst*-IRES-Cre (*Sst^tm2.1(cre)Zjh^*/J, RRID:IMSR_JAX:013044); INTACT (B6;129-*Gt(ROSA)26Sor^tm5(CAG-Sun1/sfGFP)Nat^*/J, RRID:IMSR_JAX:021039, LSL-Sun1-GFP); RiboTag (B6J.129(Cg)-*Rpl22^tm1.1Psam^*/SjJ, RRID:IMSR_JAX:029977); *Pvalb*-2A-Cre (B6.Cg-*Pvalb^tm1.1(cre)Aibs^*/J, RRID:IMSR_JAX:012358); MeCP2 Ser421Ala KI (*Mecp2^tm1^*^.1Meg^, RRID:MGI:5302547)^17^. To generate PV/ or SST/INTACT mice, homozygous *Pvalb*-IRES-Cre or *Sst*-IRES-Cre males were bred with homozygous INTACT females to create compound heterozygous offspring, which were used in all subsequent experiments. Unless explicitly stated, all experiments used adult (P60-210) male and/or female mice that were heterozygous (HET) for both the *Pvalb*-Cre or *Sst*-Cre and INTACT alleles. To generate PV/RiboTag mice on the *Mecp2* Ser421Ala KI background, we crossed female *Mecp2* Ser421Ala/*Mecp2* WT HET mice to *Pvalb-*2A-Cre mice and then crossed the offspring to one another to generate MeCP2 Ser421Ala heterozygous;*Pvalb*-2A-Cre homozygous females. These mice were crossed to homozygous RiboTag male mice, and all experiments were conducted using the adult MeCP2 KI and WT hemizygous male littermates, all of which were heterozygous for the *Pvalb*-2A-Cre and Ribotag alleles.

### Open field locomotor activity and AMPH administration

Mice were moved into the open-field testing room 24hrs before each open field trial. Each day mice were habituated to the open field for 1hr to establish baseline locomotor activity. On day one, after habituation, mice received a mock injection and were returned to the open field. Locomotor activity was monitored as horizontal distance traveled (cm). To study acute responses to AMPH, on day 2, either saline (as a vehicle control) or 3mg/kg AMPH was administered (i.p.) and mice were returned immediately to the open field for 35min (RNA-seq timepoint 1, snRNA-seq), 60min (ATAC-seq), or 3hr (RNA-seq timepoint 2). To study chronic responses to AMPH, mice were habituated as above and then given either saline or 3mg/kg AMPH once each day for days 2-8 in the open field with their locomotor activity recorded for 90min post-injection. Mice were removed from the open field and rapidly sacrificed. The NAc was dissected and flash frozen in chilled 2-methyl butane, then stored at −80°C until nuclear isolation. Tissue used for the single nuclear RNA-seq study was processed fresh and moved immediately into the nuclear isolation protocol.

### INTACT nuclear isolation

We used a variation of the published INTACT protocol^19^. For each experiment, the two NAc samples from each single mouse were thawed in ice-cold homogenization buffer (0.25M sucrose, 25mM KCl, 5mM MgCl2, 20mM Tricine-KOH). The tissue was minced with a razor blade and dounce homogenized using a loose pestle in 1.5mL of homogenization buffer supplemented with 1mM DTT, 0.15mM spermine, 0.5mM spermidine, 172g/L kynurenic acid, and EDTA-free protease inhibitor. A 5% IGEPAL-630 solution was added to bring the homogenate to 0.3% IGEPAL-630, and the homogenate was further dounced with eight strokes of the tight pestle. When purifying RNA, RNaseOUT was added to all buffers at 60U/mL. When isolating nuclei for ATAC-seq, sodium butyrate was added to all buffers at a final concentration of 5mM. The sample was filtered through a 40μm strainer, mixed with 1.5mL of Working Solution (1:5 150mM KCl, 30mM MgCl2, 120mM Tricine-KOH, pH 7.8 Diluent and Optiprep Density Gradient Medium), underlaid with a gradient of 30% and 40% iodixanol, and centrifuged at 10,000xg for 18min on an Sw41Ti rotor in a swinging bucket centrifuge at 4°C. Nuclei were collected at the 30%-40% interface and pre-cleared by incubating with 20μL of protein G magnetic Dynabeads for 10min. After removing the beads with a magnet, the mixture was diluted with wash buffer (homogenization buffer plus 0.4% IGEPAL-630) and incubated with 10μL of 0.2mg/mL rabbit monoclonal anti-GFP antibody (Thermo Fisher Scientific Cat# G10362, RRID:AB_2536526) for 30min. 60μL of Dynabeads were added, and the mixture was incubated for an additional 25 minutes. To increase yield, the bead-nuclei mixture was placed on a magnet for 30sec to 1min, completely resuspended by inversion, and placed back on the magnet. This was repeated 5-7 times. Samples were then re-placed on the magnet for 5min. 1mL of supernatant was removed as the Unbound Nuclear Fraction (UF) and placed on ice. The remaining beads were washed 3 times in 1mL of wash buffer followed by one wash in 6mL wash buffer. The 6mL was then sequentially applied to the magnet until all beads had been isolated. The final bead mixture was resuspended in 100µL wash buffer for downstream applications. All steps were performed on ice or in the cold room, and all incubations were carried out using an end-to-end rotator. 10µL of suspended UF nuclei were counted on a hemocytometer and 5000 UF nuclei were removed for use in downstream applications as an approximate reference sample to the number of PV+ nuclei harvested per mouse (calculated range 3000-5000/mouse).

### Immunostaining

Mice were anesthetized using isoflurane in a bell jar, and transcardially perfused using chilled 4% PFA in PBS. Brains were post fixed overnight at 4°C in 4% PFA in PBS, and subsequently sucrose protected in 30% Sucrose + PBS and sectioned coronally on a freezing microtome at 30µm. For IHC, sections were incubated for 60min in blocking buffer (PBS + 0.3% Triton X-100 and 10% Normal Goat Serum. Sections were then incubated overnight at 4°C in blocking buffer and primary antibody. The following primary antibodies were used: Rabbit-α-GFP (1:200; ThermoFisher G10362; RRID RRID:AB_2536526), Guinea Pig α-Parvalbumin (1:1000; Synaptic Systems, 195 004RRID:AB_2156476), Rabbit α-c-Fos (1:750, Millipore PC38, Ab-5 RRID:AB_2314421). After primary antibody incubation and washing 3 times with PBS, sections were incubated with Cy2, Cy3, or Cy5-conjugated secondary antibodies from Jackson Immunoresearch. Nuclei were counterstained with 1nM DAPI solution before mounting with Prolong Gold.

### Fluorescent in situ hybridization (FISH)

We performed RNAscope FISH (ACD) according to the manufacturer’s instructions to validate cell-type specific gene expression in the NAc and to validate quantification of differentially expressed genes detected by sequencing. Brains were harvested and flash-frozen in an isopentane/dry ice bath and embedded in Optimal Cutting Temperature (OCT) medium. 20µm coronal sections were cut on a cryostat and mounted on Superfrost Plus slides. We used the following probes (Advanced Cell Diagnostics): Mm-*Pvalb* (Cat no. 421931), Mm-*Pvalb*-C3 (Cat no. 421931-C3), Mm-*Sst* (Cat No. 404631), Mm-*Ppp1r1b*(Cat no. 405901), Mm-*Cntnap4* (Cat no. 498571), Mm-*Acan*-C2 (Cat no.439101), Mm-*Gapdh*-No-X-Hs (Cat no. 442871), Mm-*Fos*-C2 (Cat no. 316921-C2), Mm-*Adarb2* (Cat no. 519971), and Probe Diluent (Cat no. 30041). Slides were counterstained with DAPI and coverslipped using ProLong Gold mounting medium. RNAscope fluorescent signal was imaged at 63X on a Leica SP8 confocal microscope and quantified using Fiji/ImageJ. At least 25 cells per group were quantified (across at least 2 slices per animal, 3-4 animals per genotype or treatment). Seven 0.5µm z-steps centered on the largest diameter DAPI signal were collapsed into a sum projection for each cell. Background fluorescence for each channel was calculated from 4 sample ROIs and subtracted from the image. A ROI was then drawn around the DAPI signal, and the integrated density measured for each probe. In a subset of images, the average integrated density of single transcripts for each probe was measured across at least 80 9-pixel ROIs surrounding single spots. To calculate the number of transcripts per cell, the integrated density of each cell was divided by the single transcript value for that probe.

### nucRNA-seq and Analysis

RNA isolation was performed using the RNaqeuous Micro kit (Thermo Fisher) according to the manufacturer’s instructions, excepting that DNase digestion was not performed at this step, as the downstream library preparation included a DNase step. RNA was eluted in 15µL elution buffer from which 4µL was used for qRT-PCR gene enrichment validation. For library preparation, all samples were processed using the Ovation SoLo RNA-Seq kit (NuGEN Technologies) according to the manufacturer’s instructions. For the PCR amplification step, amplification cycles were determined individually for each sample according to the manufacturer’s instructions. Library concentration was assessed with the Qubit 2.0 fluorometer and checked for quality and fragment size on an Aligent Tapestation 2200 using a D1000 HS Tape. Samples were run on the Illumina HiSeq 4000 using a 50 base-pair, single-end read protocol by the Duke University sequencing core facility. Due to instrument retiring, the SST 3hr RNAseq timepoint and chronic AMPH ATAC-seq experiments were sequenced on an Illumina Novaseq6000 using a 50-base pair, paired-end, S-prime read protocol. FASTQ-formatted data files were processed using the Trimmomatic toolkit v0.38 to trim low-quality bases and Illumina sequencing adapters from the 3′ end of the reads. Only reads that were 32nt or longer after trimming were kept for further analysis. Reads were mapped to the Gencode annotation GRCm38v72 version of the mouse genome and transcriptome using the STAR RNA-seq alignment tool. Gene counts were compiled using the HTSeq tool. For this analysis, we used the standard method of only counting reads that mapped to known exons and reads with mapq > 30 were used for the differential expression analysis. Prior to differential expression analysis, genes with a counts per million (CPM) <1 for any sample were excluded. Normalization and differential expression analyses were carried out using the DESeq2 Bioconductor package with the R statistical programming environment while accounting for batch, treatment, and PCR-bottlenecking effects. The false discovery rate was calculated to control for multiple hypothesis testing. In the case of cell-type specific genes, genes with FDR<0.05 were considered significantly differentially expressed. For cell-type specific stimulus-regulated genes, an FDR<0.1 was used. GO Analysis was conducted using the Database for Annotation, Visualization and Integrated Discovery (DAVID), with enriched Biological Function categories used for analysis.

### Q-RT-PCR

4µL of isolated RNA was primed with Oligo-dT and synthesized into cDNA by Superscript II. Quantitative SYBR green PCR was performed on a QuantStudio realtime PCR machine using previously validated exon-skipping primers to validate enrichment and activity-dependent gene induction: *Pvalb*: F – CTTTGCTGCTGCAGACTCCT, R-CTGAGGAGAAGCCCTTCAGA; *Sst*: F-CCCAGACTCCGTCAGTTTCT, R-CCTCATCTCGTCCTGCTC; *Gapdh*: F-CATGGCCTTCCGTGTTCCT, R-TGATGTCATCATACTTGGCAGGTT; *Fos*: F-TTTATCCCCACGGTGACAGC, R-CTGCTCTACTTTGCCCCTTCT*; Bdnf* (exon IV): F-CGCCATGCAATTTCCACTATCAATAA, R-GCCTTCATGCAACCGAAGTATG.

### ATAC-seq

Omni-ATAC-seq was done as previously described^63^ with the sole modification of an adjusted Tn5 volume (0.5µL/sample) to avoid over transposition in a low number of input nuclei. Briefly, 5000 UF nuclei or the entirety of the bead-bound immunoprecipitated fraction nuclei (IP) were resuspended in cold RSB buffer prior to the beginning of the Omni-ATAC procedure. Libraries were made as described with the added inclusion of a 1:1 volume library cleanup with Ampure XP beads. The quality of the raw reads was determined using FastQC v0.11.2. All adaptors and reads with quality < 30 were trimmed using cutadapt v1.8.3. The reads were then aligned using bowtie2 v2.3.4.3 against the Gencode annotation GRCm38v72 reference genome with no more than 1 mismatch. The aligned reads were sorted and we filtered out unmapped reads using bedtools v2.25.0. Samples with a fraction of reads in peaks (FRIP) <0.125 or with mapped reads <4X the mean mapped reads were excluded. Peaks were called by converting to a bed file using bedtools, filtering duplicates, and calling broad and narrow peaks using MACS2 v 2.1.2 (parameters: --nomodel --shift 37 --ext 73) as suggested for ATAC peaks. The overhanging peak ends were clipped off the ends of chromosomes using KentUtils bedClipv302.The peaks for all samples in each comparison were merged (# peaks) to create a master peak file for a basis of comparison. Further analysis was carried out in R 3.4.4. A peak count matrix was creating using Rsubreadv1.28.1 to read in the ATAC counts from the merged peak regions. Prior to differential expression analysis, peaks with a CPM<1 in fewer than 3 samples were filtered out. DESeq2 v1.18.1 was then used to perform the differential peak analysis using an adjusted p-value threshold of p<0.05 for significance. For visualization these peaks were converted into bigwigs using deepTools bamCoverage at 10bpresolution (parameters: -- normalizeUsing RPKM). Spearman correlation of the of the DESeq2 variance stabilized peak counts from each sample was computed and plotted using the stats v 3.6.2 package and corrplot v 0.84 package in R respectively.

### Fluorescence activated nuclear sorting and single nucleus sequencing (FANS-snSeq)

Fresh NAc from PV/INTACT mice (n=3 Saline, n=4 AMPH) were isolated, pooled by treatment and homogenized in homogenization buffer. PV+ nuclei were isolated according to the Isolation protocol above with the exception that samples were underlaid with only a 30% iodixanol solution – allowing for the generation of a nuclear pellet after centrifugation. Nuclei were then re-suspended, washed in homogenization buffer, and incubated with MULTI-seq lipid-modified oligos (LMOs) 5 or 6 to barcode nuclei from either the saline or AMPH condition respectively^42^. LMOs were added at a ratio of 10:1 oligo barcode to molecular (lipid) anchor and DAPI. The saline and AMPH treated nuclei were then pooled for FANS, loading onto the 10X chromium controller, and sequencing to avoid batch effects. PV+ nuclei were isolated using Fluorescent-Activated Cell Sorting (FACS) gating on double positivity for DAPI and GFP and sorted into homogenization buffer. Flow cytometry was performed and nuclei were sorted directly into a plate for 10X Genomics snRNA-Seq. 10X Genomics 3’ Gene Expression library (v3 chemistry) was sequenced on Illumina Nextseq 550 in mid output mode. Raw BCL files were converted to fastqs using CellRanger v3.0.2 mkfastq. Fastq files were then aligned to the mm10-3.0.0_pre-mRNA reference transcriptome and a count matrix was generated using CellRanger v3.0.2. This count matrix was used as the input to Seurat (v3.1.5) for downstream analysis.

### snRNA-seq Analysis

snRNA-seq analysis was performed in R using Seurat v3.1.5. Cell Ranger count matrix of 2125 nuclei was read into Seurat using the Read10X command. Seurat object was created using raw cell counts with baseline criteria of min.cells = 3 and min.features = 200. For initial filtration for presumed doublets and low-read nuclei, the count matrix was filtered for nuclei containing more than 3000 and fewer than 15000 molecules detected (nCount_RNA), yielding a total of 1687 nuclei with an average of 2767 detected genes per cell and 6221 counts per cell detected. Nuclei of interest were subsequently filtered based on detectable expression of *Pvalb* transcripts >0.5 using the subset command, yielding 787 nuclei for subsequent analyses. Data was log normalized using the NormalizeData command and highly variable genes were identified using the FindVariableFeatures command (selection.method=‘vst’, nfeatures = 2,000). We then performed Principal Component Analysis using RunPCA to compute 20 components followed by dimensionality reduction with Uniform Manifold Approximation and Projection (UMAP) via the RunUMAP command using the top 2000 genes with the highest variance. A shared nearest neighbor plot was generated using FindNeighbors integrating 4 dimensions and cell clustered using the FindClusters command with a resolution=0.5. We observed a significant inflection point in variance explained after PC4 and thus proceeded with the inclusion of these 4 PCs in the generation of our UMAP. This number of dimensions was chosen for maximal variance integration while minimizing the propensity for over-clustering within an otherwise relatively homogeneous cell population. For between sub-type comparisons, identities were created based on minimum expression criteria of *Pvalb* and *Adarb2* and compared using Wilcoxon Rank Sum test via the FindMarkers command.

### Transcription factor motif analysis

We searched regions around AMPH-induced, differentially expressed (DE) genes for transcription factor binding motifs in a cell-type specific manner. We considered genes upregulated in PV+ interneurons, SST+ interneurons, or the unbound fraction (UF), which is enriched for SPNs. bedtools v2.27.1 was used to find peaks that were differentially accessible (DA) in each cell type and were either within the gene bodies of DE genes, their promoters, or distal regions within 50kb of the DE gene. Once each peak set was created, the peaks were expanded from the center to create a 500bp region for motif analysis. Motif enrichment analysis was carried out in Homer v4.10.4 separately for promoter regions and non-promoter regions (both within-gene and gene-distal regions). For the promoter motif enrichment, random peaks were chosen in the promoters of nonDE genes and used as background. For non-promoter peaks, the Homer-standard genomic regions matched for GC% were used as a background. Each significantly enriched motif (q < 0.05) was mapped to its transcription factor gene/gene families and that transcription factor was then assessed for its relative gene expression in each cell type. Transcription factors that had enriched motifs and also showed differential expression in a specific cell-type transcript were considered putative regulators of the AMPH response in that cell type.

### RiboTag purification of cell-type specific translating RNA

Ribotag purification of translating RNAs was performed with minor variations from the published protocol^51^. Briefly, NAc tissue was rapidly dissected and Dounce homogenized in a tissue weight to buffer volume ratio of 5% (0.6-1mL) of homogenization buffer (HB-S, 50mM Tris, 100mM KCl, 12mM MgCl2, 1% NP-40) supplemented with 200U/mL RNAaseOUT, 1X Protease inhibitor, 1mM DTT, 100ug/mL Cyclohexamide, and 1mg/mL Heparin. Homogenate was spun down at 10,000 RPM and the supernatant was moved to the immunoprecipitation step after removal of 80µL for input vs IP comparisons. To prepare the antibody-bead mixture, Dynabeads were first washed in 1X PBS before 10µL Rabbit α-HA tag antibody (Abcam, #9110, RRID:AB_307019) was added to 300µL PBS+60µL Dynabeads and rotated at 4°C for 4hr. For immunoprecipitation, the remaining supernatant was subsequently combined with α-HA antibody-bead mixture and rotated overnight at 4°C. Beads were isolated, washed 3 times in high-salt buffer, and lysed in Qiagen lysis buffer RLT for elution. RNA was subsequently extracted using the Qiagen RNeasy Mini Kit. SMART-Seq™ v4 Ultra™ Low Input RNA Kit (Takara Bio #634888) was used to convert RNA to cDNA, which was then amplified and sequenced using an Illumina Hi-Seq 2500.

### Statistical analyses

Unless otherwise indicated, graphs show mean and SEM with individual points shown. For comparisons of averages, data were tested for normality using the Shapiro-Wilk (SW) test. For multiple-comparison tests of locomotor activity in response to treatment, rmANOVA was performed using PRISM (GraphPad) and Bonferroni-corrected pairwise tests were used *post hoc* to correct for multiple comparisons. P values <0.05 were considered to be significant. FDR<0.05 for Cell-type-specific genes, FDR<0.1 for AMPH/Genotype regulated nuclear or ribosomal RNAseq, and FDR<0.05 for ATAC-seq data were considered significant.

## Data Availability

RNA and ATAC sequencing data that support the findings of this study will be deposited at GEO immediately following submission (accession number pending). All other primary data are stored on a secure server at Duke University School of Medicine and are available from the corresponding authors upon request.

## Code Availability

Full coding implementation of all analysis tools can be found in complete alignment/analysis pipelines available at https://github.com/WestLabDuke/Psychostimulant-NAcInterneuron.

## Acknowledgments

We thank Xiaoting Wang, Alexias Safi, and Greg Crawford for assistance with experiments. The Duke University School of Medicine Sequencing and Genomic Technologies Shared Resource provided sequencing services and the Duke University Mouse Behavioral and Neuroendocrine Analysis Core Facility provided equipment and support for the mouse behavioral studies. FANS was performed in the Immunology Unit of the Regional Biocontainment Laboratory at Duke, which receives support from NIH grant UC6-AI058607. This work was supported by NIH grants R01DA047115 and R33DA041878 (A.E.W.).

## Author contributions

D.A.G. and A.E.W. conceived of and designed the study. D.A.G., M.M., F.L., M.F.H., S.A.Y., and L.C.B. acquired and analyzed data. D.A.G. and A.E.W. wrote the paper with feedback from other authors. All authors read and approved the submitted version of the study.

## Materials and Correspondence

All correspondence and requests for material should be directed to Anne West, 311 Research Drive, Bryan Research 301D, Duke University Medical Center Box 3209, Durham, NC 27710, USA., phone 919-681-1909.

---

**Figure S1:**
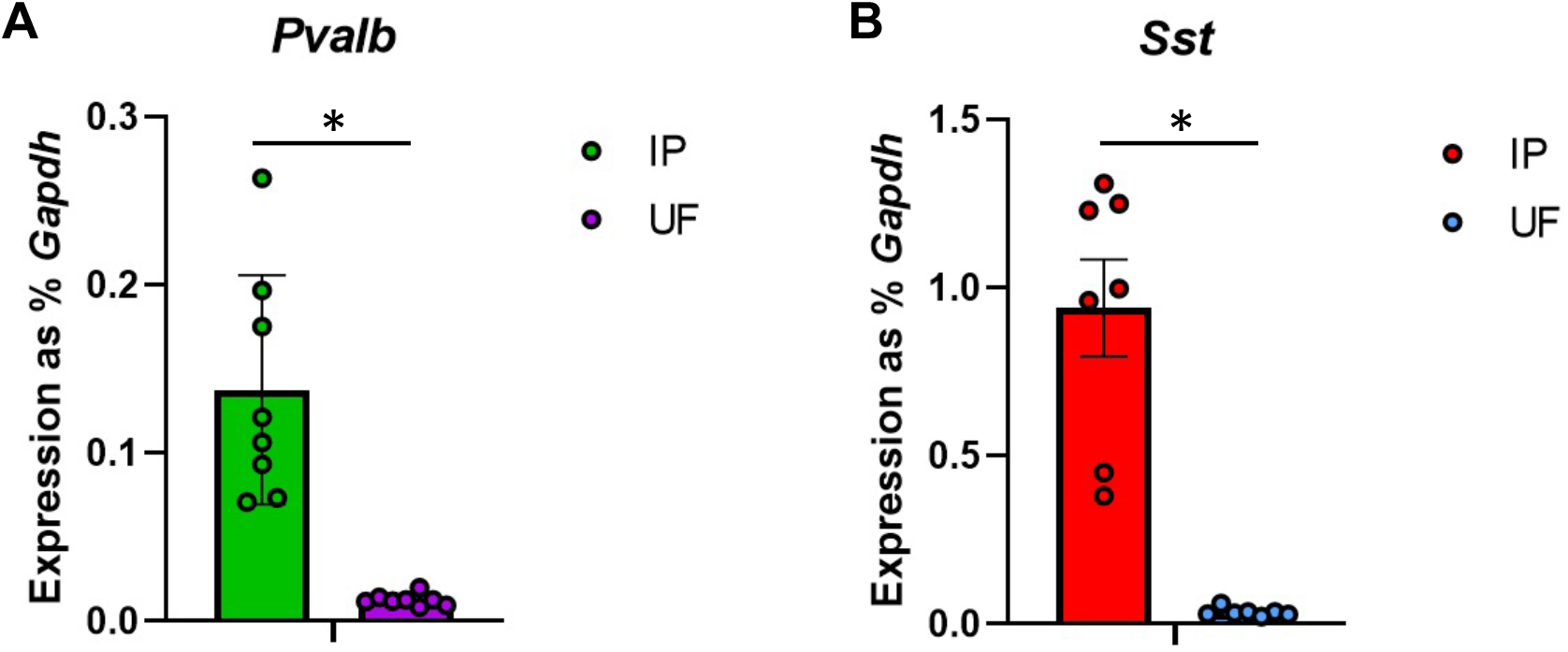
Q-RT-PCR validation of interneuron marker gene enrichment in INTACT IP fractions. Prior to the NucRNA-seq in Figure 1, portions of harvested nucRNA from each IP and UF were processed for Q-RT-PCR for *Pvalb* (**A)** or *Sst* (**B**). All samples were normalized to expression of the housekeeping gene *Gapdh* in the same sample. N=8 *Pvalb-Cre* and 7 Sst-Cre. Paired t-test, two tailed, p=.0001 (*Pvalb-*Cre), p = .0007 (*Sst-*Cre). Bars show mean and error bars indicate SEM.

**Figure S2:**
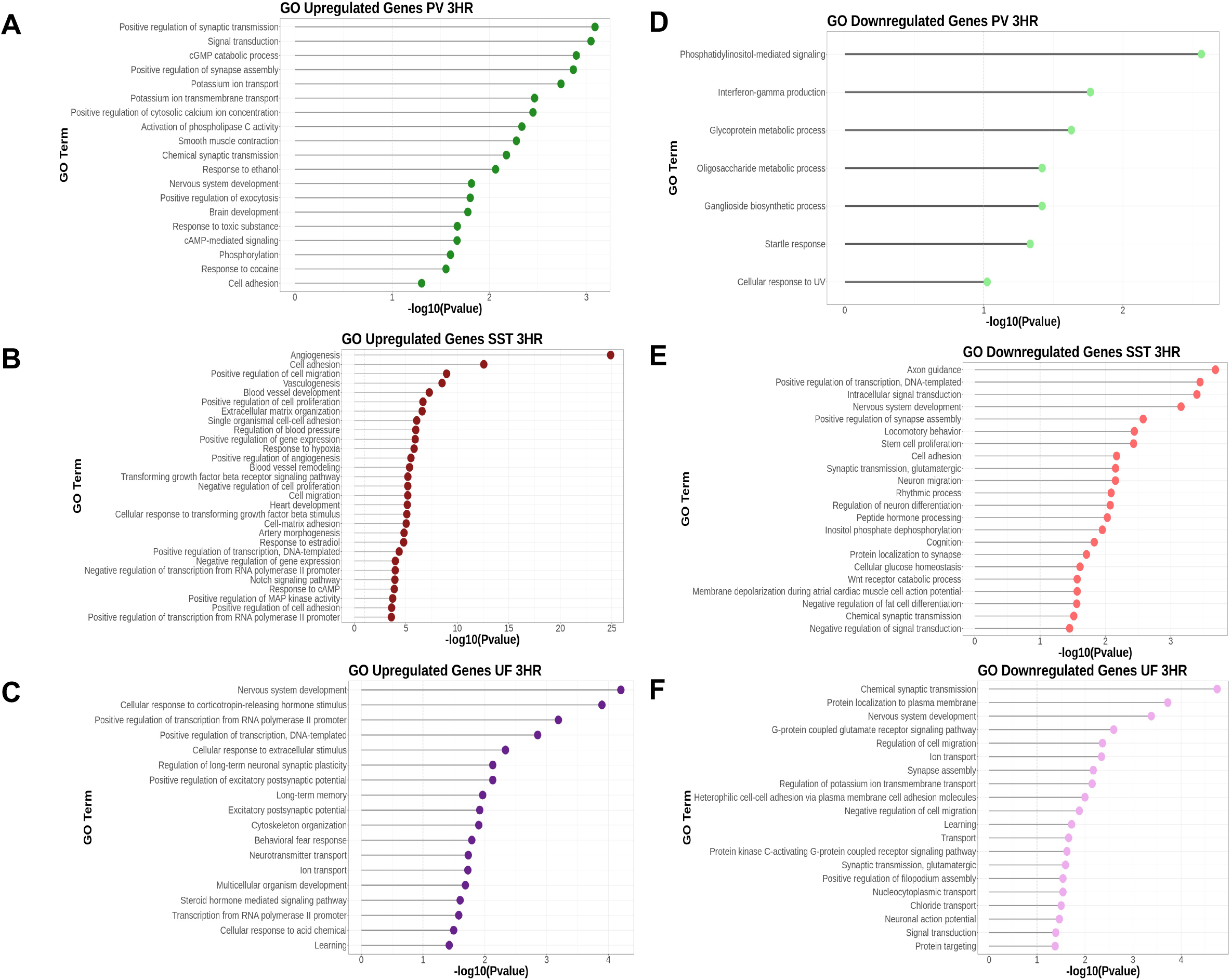
Gene Ontology Analysis of AMPH-regulated Delayed Primary and Secondary Response Gene Programs. **A-C)** DAVID Gene Ontology (GO) analysis of genes induced by AMPH at 3hr post-administration. Shown are enriched Biological Function (BF) GO terms for genes upregulated by AMPH in each population of nuclei *Pvalb-*Cre IP, Green (A); *Sst*-Cre IP, red (B); Combined UF (C), purple; *p<0.05. **D-F)** DAVID Gene Ontology (GO) analysis of genes downregulated by AMPH at 3hr post-administration. Shown are enriched Biological Function (BF) GO terms for genes downregulated by AMPH in each population of nuclei *Pvalb-*Cre IP, Green (D); *Sst*-Cre IP, red (E); Combined UF (F), purple; *p<0.05.

**Figure S3:**
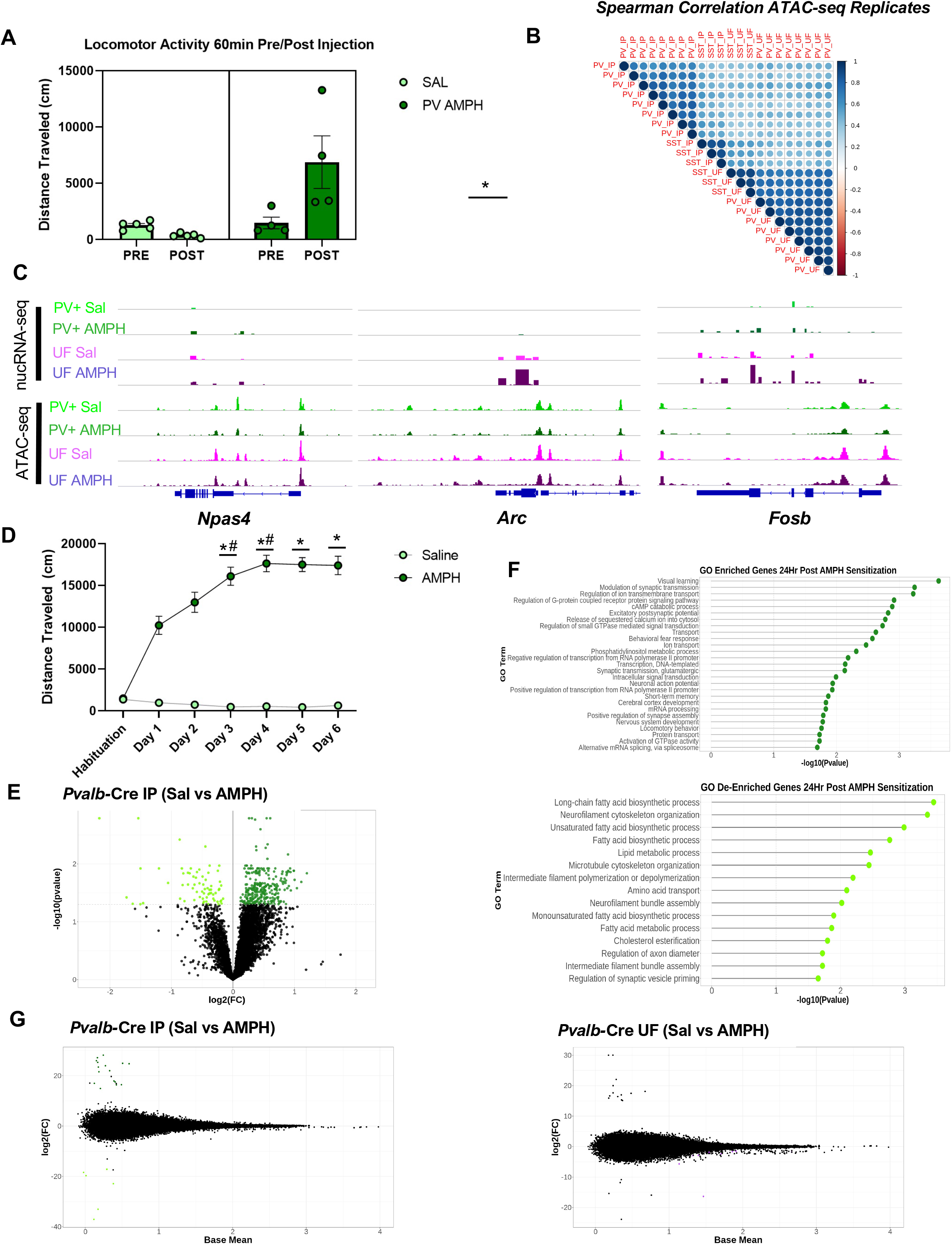
**A)** Locomotor activity 60 min before (Pre) and 60 min after (Post) acute i.p. injection of saline (light green) or 3mg/kg AMPH (dark green). *Pvalb-*Cre n=4/treatment condition; Two-way ANOVA, *Pvalb-*Cre F (1, 7) = 13.80, p=0.0075), Bonferroni’s post-hoc PV AMPH Pre vs Post p=0.0073, Error bars indicate SEM. **B)** Spearman Correlogram of variance-stabilized counts for each ATAC-seq sample (*Pvalb-*Cre IP, *Sst*-Cre IP, Combined UF). **C)** Representative nucRNA-seq tracks of AMPH-induced PRGs *Arc, Npas4,* and *Fosb* with associated ATAC-seq tracks of chromatin accessibility surrounding each gene. **D)** Post-Injection locomotor activity on each of 7 days in open field chamber. Day 0 was habituation to chamber with a mock i.p. injection, followed by daily i.p. injections of either Sal or 3mg/kg AMPH on days 1-6. ATAC-seq was performed on nuclei harvested from dissected NAc of individual mice 24hr after Day 6 injection; Locomotor sensitization is observed in mice receiving AMPH. *Pvalb-*Cre n=4/treatment condition. Mixed Effects Model, F (6, 123) = 79.67 p<.0001, Bonferroni post-hoc *p<0.01 vs Day 1; #p<0.01 vs Day 2, Error bars indicate SEM **E)** Volcano plots of differential gene expression in immunoprecipitated PV+ cells in *Pvalb*-Cre IP (green) at 24hr following daily injections of either Sal or AMPH as described above *FDR<0.05. **F)** DAVID Gene Ontology (GO) analysis of differentially expressed genes at 24hr post repeated Sal/AMPH administration. Shown are enriched Biological Function (BF) GO terms for genes upregulated (dark green) or downregulated (light green) in the immunoprecipitated population of *Pvalb-*Cre nuclei, *p<0.05 **G)** MA plots of AMPH-induced differential chromatin accessibility in each population of isolated nuclei 24-hrs after 6 days of repeated treatment with either Sal or AMPH using DeSeq2 *FDR<.05; *Pvalb-*Cre IP n=4 AMPH, n=3 Sal, *Pvalb-*Cre UF n=4/treatment condition.

**Figure S4:**
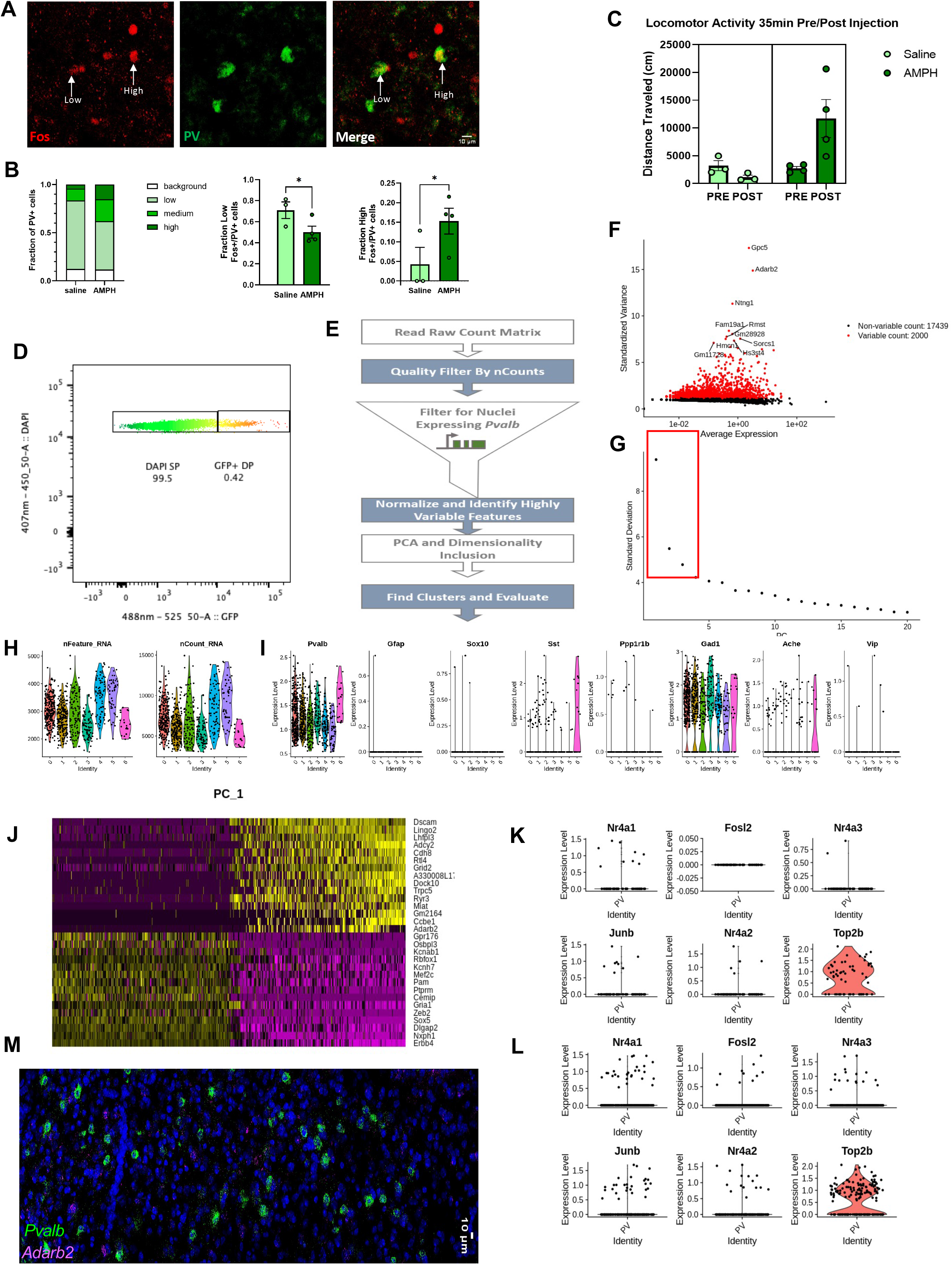
**A)** Representative images of NAc slices immunohistochemically stained with antibodies against Fos and PV 2 Hrs following acute injection of 3mg/kg AMPH. Example PV+ cells binned into low and high Fos immunofluorescence categories are indicated with labeled arrows. **B)** Quantified relative frequency of Fos IHC fluorescence intensity in immunohistochemically positive PV cells of the NAc; Cells binned into Background (0-1.5x), Low (1.5-3x), Medium (3-4.5x), and High (4.5x+) groups based on Fos fluorescence intensity relative to average background fluorescence intensity in the same channel. Per bin one-tailed t-test, *p<0.05 Error bars indicate SEM. **C)** Locomotor activity 60 min before (Pre) and 35 min after (Post) i.p. injection of saline (light green) or AMPH (dark green) for *Pvalb*-Cre mice pooled and used for FANS. *Pvalb-*Cre n=3 LMO 5/SAL, n=4 LMO6/AMPH Two-way rmANOVA, F (1, 5) = 6.402, TimeXTreatment p=0.0525, Bonferroni post-hoc p=0.0513 **D)** Fluorescence intensity plot showing fluorescence intensity at 407nm and 488nm amongst pooled *Pvalb-*Cre NAc nuclei submitted for FANS. Gating criteria/region for DAPI/GFP double-positive (DP) nuclei harvested for snRNA-seq is shown with a white square, compared to DAPI single-positive (SP) nuclei. **E)** Summary flow chart of pipeline used for Seurat analysis of snRNA-seq data. **F)** Plot of genes contributing to the greatest amount of dispersion (Variance contribution vs average expression) using FindVariableGenes function in Seurat 3.1.5 **G)** Elbow plot of variance explained by each Principal Component (PC) using the RunPCA function in Seurat 3.0; PCs/dims used for downstream UMAP generation are indicated by a red box. **H)** Violin plot of Features/Genes as well as Counts/Molecules detected in each nucleus across the 6 UMAP projection clusters snRNA-seq following filtration based low counts and detectable *Pvalb* transcripts. **I)** Violin plots of log-normalized expression levels of NAc cell-type marker genes in nuclei across the 6 UMAP projection clusters. **J)** Heatmap of high-variance genes contributing to integrates PC1 **K-L)** Violin plots of log-normalized expression levels of various PRGs in nuclei confirmed positive for Multi-seq LMO 5 (K, Sal treated n=60) or LMO6 (L, AMPH-treated, n=187).

**Figure S5:**
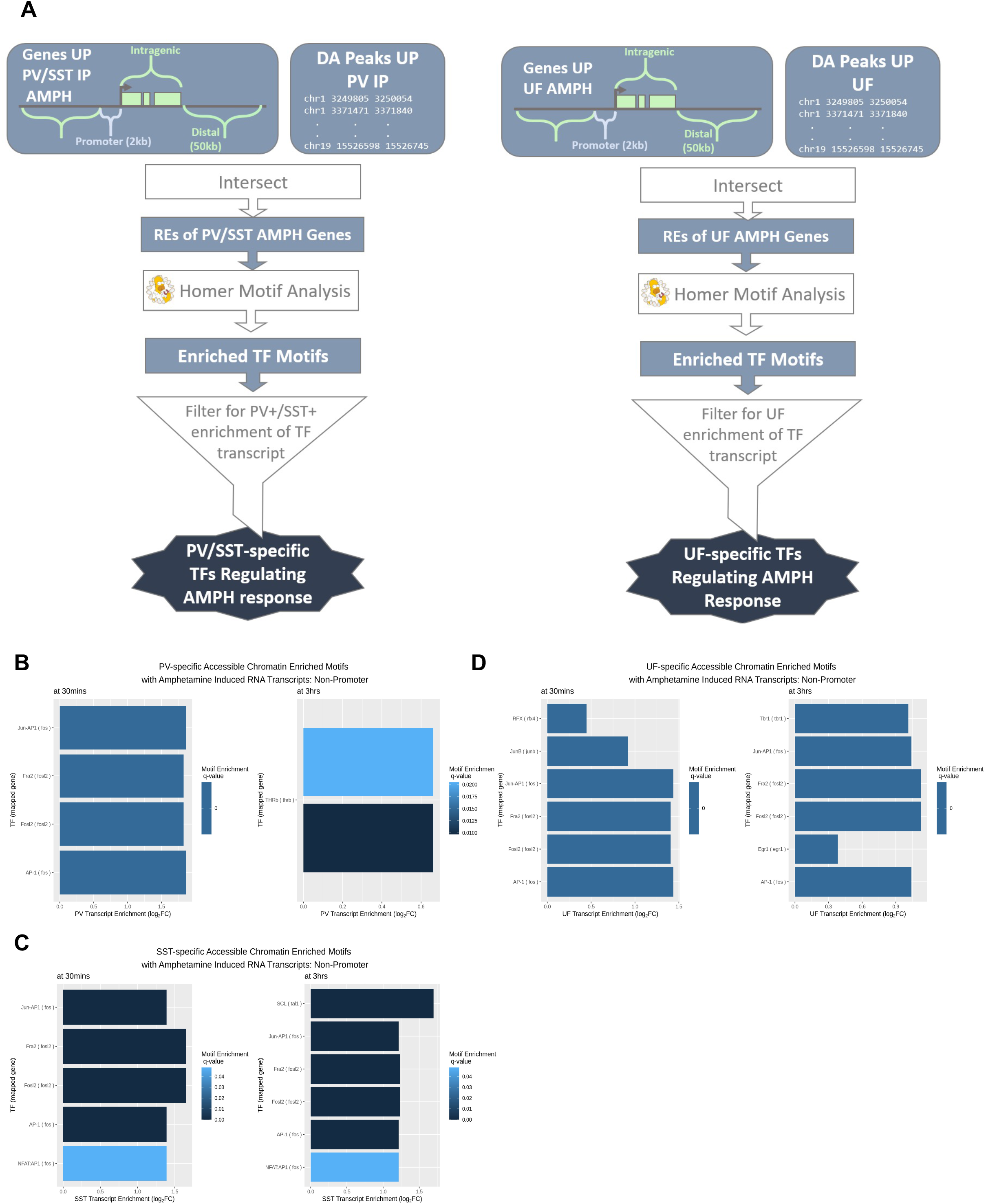
**A)** Informatic flow chart of Transcription Factor Motif enrichment and RNA-enrichment analysis pipeline in *Pvalb-*Cre/*Sst-*Cre IP and Combined UF nuclei. **B-D)** Enriched Transcription Factor Motifs as determined by HOMER at cell-type-unique differentially accessible inter- (+/− 50Kb) and intragenic chromatin regions at genes induced by AMPH at 3Hrs in each cell fraction, *q<.05; Enriched motifs are plotted against log2FC of induction by AMPH at 35min or 3hrs post AMPH of cognate RNA transcript in each isolated cell type or UF; *Pvalb*-Cre IP vs UF (B), *Sst-*Cre IP vs UF (C), Combined UF vs Combined IP (*Pvalb*-Cre+*Sst-*Cre IP).

**Figure S6:**
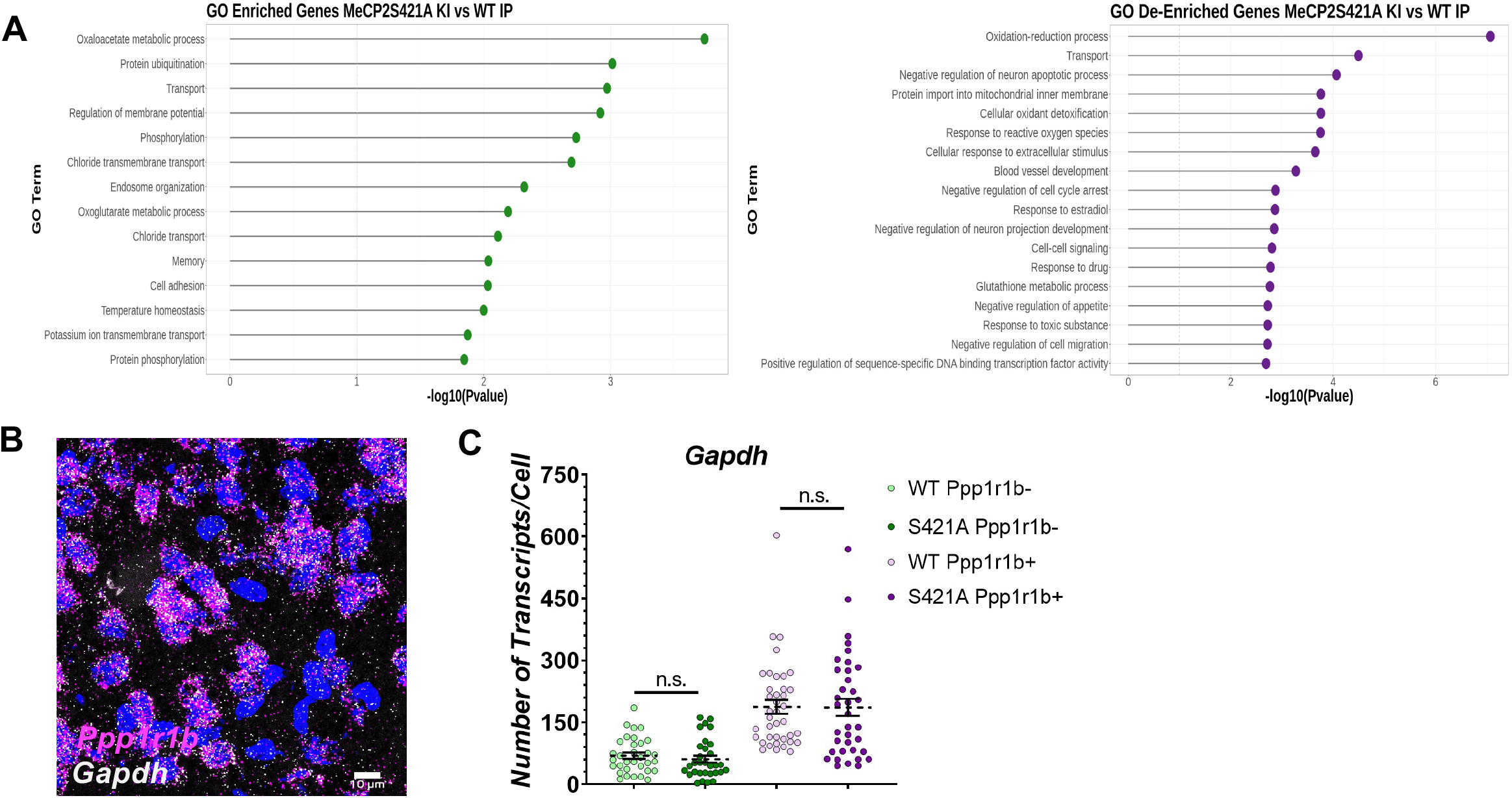
**A)** DAVID Gene Ontology (GO) analysis of differentially expressed genes between MeCP2Ser421Ala KI IP and WT IP. Shown are enriched Biological Function (BF) GO terms for genes upregulated (dark green) or downregulated (purple) in the immunoprecipitated population of *Pvalb-*Cre nuclei *p<.05 **B)** Representative smFISH image of *Ppp1r1b* and *Gapdh* fluorescent labeling **C)** Reference smFISH quantification of *Gapdh* transcript number based on punctate fluorescent signature in *Ppp1r1b+* or *Ppp1r1b-*nuclei (n=39 WT *Ppp1r1b+*, 36 KI *Ppp1r1b+* nuclei, 33 WT *Ppp1r1b-*nuclei, 31 KI *Ppp1r1b+* nuclei) Two-way ANOVA, F (1, 135) = 0.1044, p=.7471.

## List of Supplementary Tables

**Table S1-NAc PV and SST nucRNAseq**

Differentially expressed genes (DEGs) calculated comparing nuclear RNAseq data from immunoprecipitated (IP) nuclei of PV or SST neurons compared with nuclei of their respective Unbound Fractions (UF). Data are shown in TPM (transcripts per million bases mapped). Data for all genes is shown on the tabs labeled ALL.

**Table S2-Rapid AMPH nucRNAseq**

Differentially expressed genes (DEGs) 35 min after exposure to 3mg/kg AMPH (i.p.) compared against saline (SAL) as control. Nuclear RNAseq data is from immunoprecipitated (IP) nuclei of PV or SST neurons or cells from the combined Unbound Fractions (UF). Data are shown in TPM (transcripts per million bases mapped). Data for all genes is shown on the tabs labeled ALL.

**Table S3 -Delayed AMPH nucRNAseq**

Differentially expressed genes (DEGs) 3hr after exposure to 3mg/kg AMPH (i.p.) compared against saline (SAL) as control. Nuclear RNAseq data is from immunoprecipitated (IP) nuclei of PV or SST neurons or cells from the combined Unbound Fractions (UF). Data are shown in TPM (transcripts per million bases mapped). Data for all genes is shown on the tabs labeled ALL.

**Table S4-NAc PV and SST ATACseq**

Differentially open chromatin regions calculated comparing nuclear ATACseq data from immunoprecipitated (IP) nuclei of PV or SST neurons compared with nuclei of their respective Unbound Fractions (UF).

**Table S5-Rapid AMPH ATACseq**

Differentially open chromatin regions in either NAc PV neurons (IP) or nuclei of the unbound fraction (UF) 60 min after exposure to 3mg/kg AMPH (i.p.) compared against saline (SAL) as control.

**Table S6-Repeated AMPH nucRNAseq**

Differentially expressed genes (DEGs) 24hr after 6 days of exposure to 3mg/kg AMPH (i.p.) compared against saline (SAL) as control. Nuclear RNAseq data is from immunoprecipitated (IP) nuclei of PV neurons or cells from the Unbound Fraction (UF). Data are shown in TPM (transcripts per million bases mapped). Data for all genes is shown on the tabs labeled ALL.

**Table S7-Repeated AMPH ATACseq**

Differentially open chromatin regions in either NAc PV neurons (IP) or nuclei of the unbound fraction (UF) 24 hrs after 6 days exposure to 3mg/kg AMPH (i.p.) compared against saline (SAL) as control.

**Table S8-NAc PV+ snRNAseq**

Gene counts per single nucleus filtered for FACS sorted nuclei with *Pvalb* expression. Tabs show the most variable genes across all nuclei and between *Adarb2*+ and *Adarb2*-populations.

**Table S9-TF motifs in open chromatin near AMPH-regulated genes**

TF motifs identified by Homer in celltype differentially accessible regions of chromatin flanking genes that show regulation by AMPH. AMPH regulation of TF expression is from Table S3.

**Table S10-NAc PV+ RNA in MeCP2WTvsKI**

Differentially expressed genes (DEGs) in translating RNA expressed in NAc PV+ neurons of MeCP2 WT or Ser421Ala KI mice. Average values shown in FPKM. Data for all genes is shown on the tabs labeled ALL.

